# LipoGrid: A High-Throughput Multi-omics Perturbation Screen Dissects the Genetic Architecture of Lipid Metabolism

**DOI:** 10.64898/2026.08.12.744385

**Authors:** Jelle Jacobs, Paulien Van Minsel, Nina Ravoet, Emiel De Rieck, Niels Vandermeulen, Lorenzo Venturelli, Katy Vandereyken, Karen Ven, Jolien Breukers, David Wouters, Thierry Voet, Jeroen Lammertyn, Johannes V. Swinnen, Bernard Thienpont, Alejandro Sifrim

## Abstract

Lipids constitute one of the largest and most diverse classes of cellular molecules, sustaining membrane architecture, energy storage, and signaling. Consequently, their dysregulation underlies a broad spectrum of human disease. However, the genetic mechanisms governing lipid homeostasis have remained largely inaccessible, owing to the lack of approaches capable of systematically linking defined genetic perturbations to large-scale changes in cellular lipidome composition. Here we introduce LipoGrid, a spatial mass spectrometry platform that resolves the genetic architecture of lipid metabolism at single-cell resolution. LipoGrid arrays CRISPR/Cas9-perturbed cells on a micropatterned grid and sequentially captures lipidomic and gRNA identity from the same cells, complemented by single-cell RNA sequencing of matched cell populations subjected to the same perturbations. Using this approach, we quantified the relative abundance of 158 distinct lipid species across 143 target genes in a rigorously controlled experimental framework. We find that most gene knockouts produced measurable alterations in lipid composition, often affecting specific lipid classes and molecular subspecies. The screen accurately recapitulated established gene-lipid relationships, including enzyme-substrate specificities, lipid pathway regulators, and disease-associated loss-of-function phenotypes, thereby demonstrating the sensitivity and accuracy of LipoGrid. By jointly profiling transcriptomic and lipidomic responses, we further uncover compensatory feedback mechanisms that buffer the impact of genetic perturbations on the cellular lipidome. Collectively, these findings establish LipoGrid as a scalable multimodal platform for systematically mapping gene-lipid interactions and reveal the regulatory networks linking gene perturbation, transcriptional adaptation, and lipidome remodeling.

**Highlights:**

- Micropatterned single-cell growth enables spatial lipidomic perturbation screens
- LipoGrid maps 143 gene knockouts to 158 lipid species and transcriptomic states
- Perturbed lipidomes reveal compensatory feedback and lipid-class-specific uptake
- Recovers enzyme substrate specificities and disease-linked lipid signatures

## Introduction

Lipids are among the most abundant and diverse components of eukaryotic cells, fulfilling crucial functions in cell structure, energy storage and cell signaling^1,2^. Unsurprisingly, mutations in enzymes related to lipid metabolism cause congenital disorders, and altered lipid profiles are associated with tumor progression and neurodegeneration^3,4^. Despite their fundamental importance, the molecular processes that govern lipid biosynthesis and homeostasis remain incompletely understood. This knowledge gap stems in part from the vast diversity of lipid molecules and the complexity of the enzymatic pathways that determine lipid production and modification, which depend on the availability of substrates and the expression of a specific cellular enzymatic repertoire^1,5^. As a result, a comprehensive mechanistic understanding of the non-linear interplay between the expression of genes encoding lipid metabolizing enzymes (the transcriptome) and the resulting lipid composition of cells (the lipidome) is still lacking. Addressing this challenge requires the interdisciplinary development of high-throughput approaches that directly link perturbations in the molecular machinery underlying lipid biosynthesis and homeostasis to changes in cellular lipid profiles.

To address these conceptual challenges, the technological capacity to measure lipids has advanced rapidly^6^. Mass spectrometry (MS)-based lipidomics can now resolve and quantify hundreds to thousands of individual lipid species across most major lipid classes, transforming the lipidome from a handful of coarse readouts into a richly detailed molecular phenotype^6^. Most of these workflows, however, operate on bulk extracts from large numbers of cells, which renders high-throughput perturbation screening impractical^7^. Matrix-assisted laser desorption/ionization mass spectrometry imaging (MALDI-MSI) has begun to close this gap, enabling spatially resolved lipid measurements at, or approaching, single-cell resolution^8,9^. Coupling such single cell lipidomic readouts to defined genetic perturbations would, in principle, allow us to directly dissect the contribution of individual genes to the cellular lipid composition.

In this context, CRISPR-Cas9 technologies have transformed high-throughput genetic screening over the past decade by enabling guide RNA (gRNA) libraries to target hundreds to thousands of genes within a single experiment^10^. Early CRISPR screens largely assessed the impact of these perturbations on cell proliferation, viability, or marker gene expression, and more recent approaches have incorporated whole-transcriptome readouts^11^. In contrast, when assessing the effects of perturbations on lipid metabolism, studies have remained limited to the role of a few selected lipid metabolic genes in cell survival, or on changes in a limited number of lipid species. For example, Köberlin et al. combined RNAi-mediated knockdown with shotgun lipidomics to reconstruct a conserved, coregulated lipid network that modulates innate immune signaling^12^. More recently, pooled CRISPR screens incorporating lipid-related phenotypes identified CLPTM1L as a lipid scramblase essential for glycosylphosphatidylinositol biosynthesis^13^. Similarly, individual gene knockouts combined with untargeted metabolomic and lipidomic profiling have linked specific enzymes to defined lipid-metabolic phenotypes, as demonstrated for the mitochondrial folate enzyme Aldh1l2 in mouse^14^.

In parallel, dedicated single-cell mass-spectrometry platforms have begun to deliver lipid and metabolite readouts at the throughput and single-cell resolution that such screens would demand^6^. One approach captures cells on printed grids of antibody- or lectin-coated micro-capture spots and uses a convolutional neural network to locate individual cells. By performing sequential MALDI-MSI, they were able to measure lipids and N-glycans from the same cells^15^. Complementarily, HT SpaceM couples optimized small-molecule cell preparation with custom laser-etched slides to perform MALDI-MSI on a monolayer of ∼1000 cells per well. They then computationally deconvolve this signal to resolve single-cell metabolic heterogeneity, detecting around a hundred metabolite ions per experiment^16^. These new platforms establish that broad small-molecule and lipid phenotypes can now be measured or deconvolved to individual cells at scale. Beyond single-cell profiling, recent advances have enabled the integration of spatial small-molecule imaging with complementary molecular modalities on the same sample. Sequential MALDI-MSI and imaging mass cytometry (IMC) combine local lipid profiles with protein-based cell-type annotation, enabling spatial characterization of tissue microenvironments^17^. Complementarily, the Spatial Multimodal Analysis (SMA) workflow overcame the longstanding incompatibility between MALDI-MSI and spatial transcriptomics, enabling the joint measurement of spatial transcriptomes and MALDI-derived metabolite and lipid profiles from the same tissue section^18^. This matched-section transcriptome–metabolome strategy has since been extended to integrative analyses of tissue injury^19^. Together, these approaches establish that spatial lipidomic and metabolomic information can be co-registered with either protein- or RNA-based cellular identities.

Despite these advances, a platform for systematically mapping genetic perturbations to single cell lipidomic phenotypes at scale has remained lacking. Achieving this requires the integration of defined genetic perturbations with comprehensive lipidomic profiling in a high-throughput, batch-controlled framework. To address this challenge, we developed LipoGrid, a multimodal screening platform that combines pooled CRISPR perturbations with spatial single-cell lipidomics and imaging-based gRNA identification on a micropatterned cell grid, complemented by parallel single-cell transcriptomic profiling. LipoGrid builds on five key components (Fig. 1A): (1) parallelized genetic perturbation of selected target genes using CRISPR-Cas9; (2) growth of individual cells on a micropatterned grid of cell-adhesive proteins; (3) lipidomic profiling of individual cells by spatial lipidomics (MALDI-MSI); (4) multiplexed single-cell detection of gRNA expression using an imaging-based spatial transcriptomics platform (Xenium); and (5) parallel transcriptomic profiling of individual cells by scRNAseq. This design enables internally controlled measurement of how hundreds of targeted genetic perturbations reshape the cellular lipidome, while allowing the integration of lipidomic and transcriptomic responses to each perturbation.

**Figure 1.**
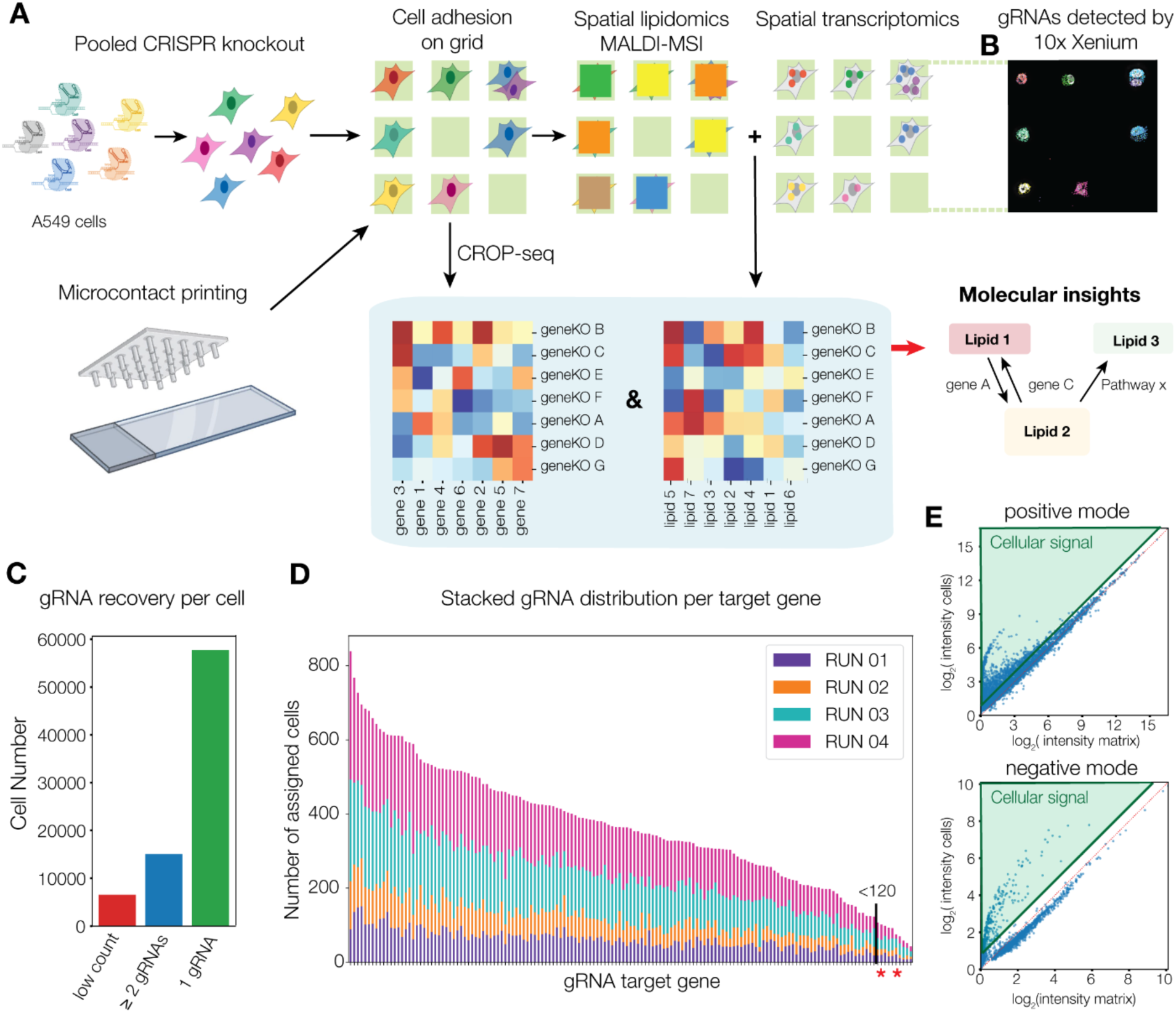
LipoGrid workflow and initial outputs. (A) Schematic overview of the LipoGrid workflow. A CROP-seq library containing 479 distinct gRNAs was delivered to A549 cells expressing inducible Cas9 by lentiviral transduction. Knockout cells were seeded onto microgrids and sequential spatial lipidomics (MALDI-MSI) followed by spatial transcriptomics (10x Xenium) was performed. In parallel single cell transcriptomic was performed on the same pool of cells. (B) Zoomed-in region of the 10x Xenium output, with each color representing a distinct gRNA. (C) Distribution of gRNA assignments across single cells. A total of 6,733 cells had no or low-confidence gRNA detected, 15,045 cells contained two or more detected gRNAs, and 57,743 cells expressed a single dominant gRNA. (D) Stacked bar plot visualizing the distribution of the different genes targeted by gRNAs in each of the 4 runs. * denotes the lethal controls SNRNP200 and PSMB3 (E) Scatterplots showing the log_2_ transformed mean intensity of each detected mass-to-charge (*m/z*) feature in positive and negative ionization modes. The x-axis represents the mean intensity measured in the surrounding matrix, and the y-axis the mean intensity measured in cells. Cell-enriched features (fold change > 1.8) are highlighted.

Applying LipoGrid to human A549 cells, we map the genetic architecture of lipid metabolism. The platform recovers expected enzyme-substrate relationships and known disease-associated lipid signatures, while the paired transcriptome reveals compensatory programs that buffer these perturbations, ranging from feedback regulation within biosynthetic pathways to induction of the corresponding lipid uptake machinery.

Together, LipoGrid provides a generalizable, cell-type- and gene-set-agnostic framework for dissecting how genetic perturbations remodel the lipidome and the transcriptional programs that maintain lipid homeostasis, offering a powerful approach for studying lipid metabolism in both physiological contexts and diseases driven by lipid dysregulation.

LipoGrid yields a controlled, paired atlas of transcriptomic and lipidomic responses spanning 143 perturbations. We provide this resource as an openly accessible database through an interactive user interface (https://sifrimlab.org/LipoGrid, see methods) that the wider lipid community can mine for their own genes and lipid species of interest. Users can retrieve the lipidomic consequences of perturbing a given enzyme, or, conversely, the genetic regulators that shape a particular lipid, thereby generating new mechanistic hypotheses.

## Results

### Developing LipoGrid

To enable pooled genetic dissection of lipid metabolism, we constructed a library of 444 gRNAs targeting 148 genes, selected for their putative role in lipid biosynthesis and homeostasis. We added 15 gRNAs targeting 5 genes as positive controls: 2 lethal genes and 3 genes encoding transcriptional regulators. For internal normalization, we also added 20 gRNAs targeting random intergenic regions (Table S1). In total, 479 gRNAs were thus designed. We optimized them both for their predicted efficiency using VBC scoring^20^ and for direct detection with custom probes for read-out on the 10x Xenium spatial transcriptomics platform (Methods). The gRNA pool was cloned into the CROP-seq^11^ vector for lentiviral particle production and transduced at a low multiplicity of infection into a clonal human epithelial cell line (A549) with a doxycycline-inducible Cas9 cassette (Methods).

Next, we developed a high-throughput method that measures lipid abundance in these cells at single cell resolution whilst preserving mRNA and gRNA readouts. We opted for Matrix-Assisted Laser Desorption/Ionization Mass Spectrometry Imaging (MALDI-MSI)^9,21^, which enables unbiased detection of abundant lipids from cells cultured on a glass slide. A spatial sampling resolution of 30 × 30 µm was chosen to allow complete acquisition of the Xenium slide within approximately 8 hours, thereby minimizing RNA degradation. Because conventional MALDI-MSI pixel sizes often encompass signal from multiple adjacent cells, we implemented, for the first time, a micropatterned grid of fibronectin patches directly on Xenium slides using microcontact printing (Figures 1A and S1A-C). The patch geometry was optimized to enable single-cell-resolved MALDI-MSI while maintaining high cell throughput, and to maximize single-cell occupancy of A549 cells while ensuring robust patterning fidelity (Methods). A patch size of 30 µm provided the best overall performance, allowing reproducible patterning while maintaining a high single-cell trapping efficiency (up to 55%). Nine days after Cas9 induction, cells were seeded onto the patterned Xenium slide, allowing knockout phenotypes to manifest before lipidomic profiling. After overnight adhesion, cells were fixed and MALDI-MSI was performed on the slide. Subsequently, cells were imaged on the 10x Genomics Xenium platform to detect expressed gRNAs (Methods). Importantly, MALDI-MSI preserved cellular morphology and RNA integrity, enabling direct linkage of each cell’s lipidomic profile to its underlying genetic perturbation (Figures 1B and S2A-C).

Finally, we performed parallel CROP-seq on the same perturbation library to capture genome-wide transcriptional responses (Methods). Integrating shared perturbation identities across the lipidomic and transcriptomic datasets linked single-cell lipid phenotypes to their corresponding transcriptional programs.

### Aligning and jointly preprocessing multimodal data

To systematically assess the effects of the selected gene perturbations on the A549 lipidome, we performed four independent LipoGrid experiments. The first two profiled lipids in positive ion mode only, whereas the latter two used sequential positive and negative ion mode acquisition to expand lipidome coverage by capturing lipid species that ionize preferentially under different conditions (Methods). Because single-cell MALDI-MSI and gRNA expression were acquired on separate imaging platforms, the datasets were spatially aligned to link each cell’s genetic perturbation with its lipidomic profile (using FOCUS^22^). This yielded a single-cell dataset comprising 79,521 single-cells, each with its *m/z* intensity profile and corresponding gRNA counts.

To assign genetic perturbations to individual cells, we analyzed the 10x Xenium custom panel data using a custom cell segmentation and gRNA assignment pipeline (Methods). We detected 459 of the 479 gRNAs, confirming robust guide detection and indicating that most genes are targeted by three independent guides. Consistent with the experimental design, 57,743 out of 79,521 cells expressed a dominant gRNA, 15,045 expressed multiple gRNAs, and 6,733 lacked a dominant gRNA assignment (Figure 1C, Methods). Cell coverage across target genes was consistent across the four runs and followed the expected distribution: a small subset of perturbations was enriched, consistent with a growth advantage, whereas some others were depleted. Depleted targets included the lethal controls *SNRNP200* and *PSMB3*, as well as several essential lipid metabolism genes, including *PISD* and *GNPAT*, which we excluded from downstream analyses. Applying a coverage threshold of more than 120 cells, exceeding that of the lethal controls, retained 143 of the 153 targeted genes for subsequent analysis (Figure 1D; Table S2). These findings are consistent with previous CROP-seq studies^11^ and with parallel CROP-seq analyses of these cells (see later), validating the robustness of the screen and demonstrating reliable gRNA detection by 10x Xenium after MALDI-MSI. Consistent with efficient editing, 86% of detected targets showed reduced expression of their own transcript in the parallel CROP-seq data, as expected from nonsense-mediated decay of frameshifted mRNAs (median log₂FC −0.15, versus 0.00 for the same genes across all other knockouts; paired Wilcoxon P = 2.3 × 10⁻^17^).

To quantify the impact of gene perturbations on the cellular lipidome, we integrated the MALDI-MSI datasets from four LipoGrid runs using a custom preprocessing pipeline^22^ (Methods), generating a unified feature matrix containing 5,472 *m/z* features in positive mode and 1,948 in negative mode. To remove background-derived signals, we used the micropatterned grid together with Xenium-defined cell coordinates to distinguish cellular regions from cell-free matrix areas (Fig. 1E). Finally, while technical variation between MALDI-MSI runs remains a major challenge for quantitative comparisons^23^, LipoGrid’s inclusion of intergenic gRNA control cells in each experiment enabled internal batch correction improving the detection of perturbation-specific lipidomic effects (Methods; Figure S3). Following quality control and feature refinement, the final dataset comprised 51,507 high-quality cells with a dominant gRNA and 1,160 positive-mode and 217 negative-mode *m/z* features for downstream analysis.

### Linking gene knockouts to changes in cellular molecular profiles

After linking single cell lipidomic profiles to defined genetic perturbations, we next asked whether these perturbations revealed structured shifts in lipidomic profiles. First, we assessed how perturbing each of the 143 target genes affected the measured *m/z* spectra. We compared the average *m/z* value of sets of cells grouped by gene knockout, contrasted to the average of all internal control cells (n=1,560). Agglomerative clustering of target genes and *m/z* ratios based on the resulting log₂ fold changes revealed clear data structures (Figure 2A). We identified seven *m/z* clusters that exhibited coordinated changes in response to specific gene knockouts (boxplots Figure S4A). Clusters 3, 4, and 5 were consistently increased by subsets of gene knockouts and decreased by others. Cluster 1 had a global decrease, while cluster 6 showed a global increase, with a subset of gene knockouts yielding more specific patterns. Finally, *m/z* values in clusters 2 and 7 responded to few or no gene knockouts (mean log₂FC -0.05 and +0.08), indicating that most features in these clusters are not affected by the targeted genes (Figures 2A and S4B-C). Together, these results indicate that most gene knockouts markedly altered a subset of the cellular *m/z* spectrum, leading to a significant increase in intensity for certain ratios and a decrease in others.

**Figure 2.**
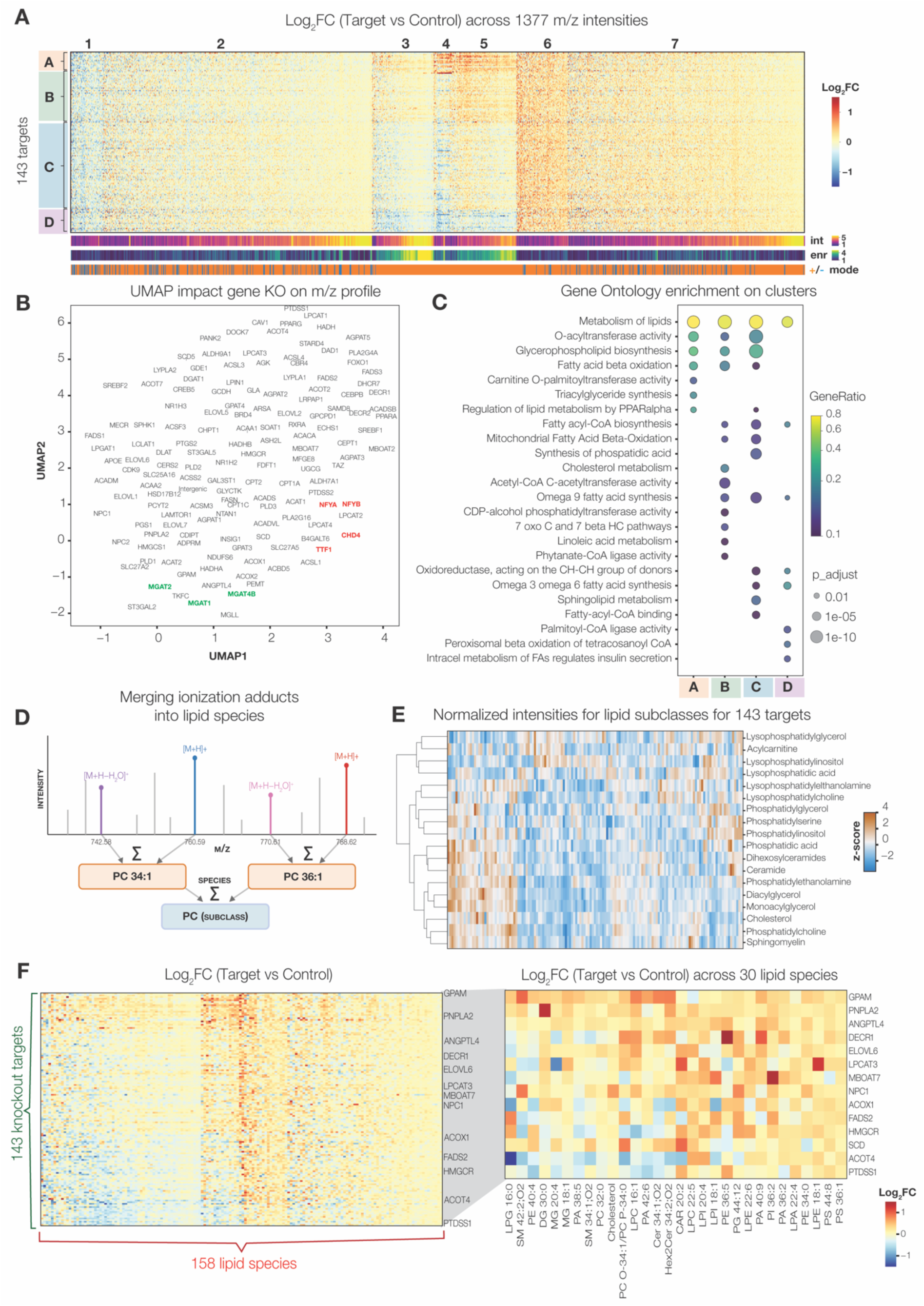
Genetic perturbations lead to significant changes in the cellular lipidome. (A) Heatmap showing the log_2_ fold change in mean intensity for each cellular enriched *m/z* feature (1,377 columns) following gene knockout relative to intergenic controls across the 143 target genes (rows). Rows and columns are ordered using agglomerative hierarchical clustering. The annotation bars beneath the heatmap indicate, for each *m/z* feature, the log_2_ transformed total mean intensity, the log_2_ fold enrichment in cells relative to the surrounding matrix, and the MALDI ionization mode (positive or negative). (B) UMAP projection of the mean *m/z* spectra (1,377 features) for each target gene knockout and the intergenic control guides. (C) Gene Ontology enrichment bubble plot showing significantly enriched terms for each of the four target gene groups defined in panel A. Bubble size represents the adjusted P value, and color indicates the overlap with the query gene set. (D) Schematic illustrating part of an *m/z* intensity profile as measured by MALDI-MSI. Signal intensities corresponding to different ionization modes of the same lipid species are summed. (E) Heatmap showing the row-wise z-score-normalized intensity profiles of each detected lipid subclass (rows) across all gene knockouts (columns). Lipid subclasses are ordered by hierarchical clustering using Ward’s linkage. (F) Heatmap showing the log_2_ fold change in mean intensity for each detected lipid specie following gene knockout relative to intergenic controls across the 143 target genes (rows). Rows and columns are ordered using agglomerative hierarchical clustering. Subset displays a selection of 30 lipid species and 14 gene knockouts.

We next assessed whether changes captured by LipoGrid hold biological meaning, by grouping the target genes according to their effects on the *m/z* intensity spectra and examining the functions of the resulting groups. Both agglomerative clustering and Uniform Manifold Approximation and Projection (UMAP) of target genes revealed a clear structure (Figures 2A-B), with co-clustering of genes encoding functionally related proteins. Examples include transcriptional regulators (NFYA, NFYB, TTF1 and CHD4) and the 3 Mannosyl-glycoprotein N-acetylglucosaminyltransferases (MGAT1, 2 and 4B) (Figure 2B, gene sets highlighted in red and green, respectively). To further test this without relying on visual inspection, we ranked all 10,153 pairs of target genes by the correlation between their lipidomic profiles. The three most similar pairs in the entire screen were ACADSB–ACADS (r = 0.82), SCD5–SCD (r = 0.81) and HADH–HADHB (r = 0.67), in each case two isozymes or two subunits of the same enzyme complex (P ≈ 10⁻⁷ that the top three pairs would all fall among the 51 same-family pairs by chance). Family membership was not by itself predictive across the whole panel, indicating that the screen recovers the closest functional relationships rather than broad sequence similarity.

To evaluate whether broader functional relationships are captured, we assessed Ontology enrichment (g:Profiler^24^) on the 4 gene sets we identified by agglomerative clustering (Figure 2A). General lipid-metabolism terms were enriched across all clusters, as expected given that the targeted genes were preselected for roles in lipid synthesis and homeostasis (Figure 2C).

However, each cluster showed specific enrichment for more specialized lipid metabolism functions. Genes in cluster A were specifically enriched for carnitine O-palmitoyltransferase activity and triacylglyceride synthesis, both governing the fate of fatty-acyl-CoA, channeling it respectively towards mitochondrial β-oxidation or storage as neutral lipid. Cluster B genes were enriched for cholesterol metabolism, acetyl-CoA C-acetyltransferase and CDP-alcohol phosphatidyl-transferase activity, representing changes in acetyl-CoA-derived membrane-lipid biosynthesis through the mevalonate pathway that feeds cholesterol synthesis, alongside phospholipid head-group assembly. Genes in cluster C were involved in many metabolic processes and specifically enriched for synthesis of phosphatidic acid (PA), sphingolipid metabolism and fatty-acyl-CoA binding, all terms linked to acyl-CoA-dependent synthesis of complex lipids, radiating from PA as the central branchpoint of glycerolipid and, in turn, sphingolipid production. Finally, genes in cluster D were uniquely enriched for palmitoyl-CoA ligase activity, peroxisomal beta oxidation of tetracosanoyl CoA and metabolic regulation of insulin secretion, the latter ontology term sharing acyl-CoA-handling genes. These terms reflect a shared role in fatty-acid activation and disposal.

These findings indicate that gene knockouts producing similar changes in cellular *m/z* intensity profiles tend to share enzymatic functions and participate in related pathways, suggesting that LipoGrid captures functionally related protein activities.

### Gene knockouts reshape cellular lipid composition

Having confirmed that LipoGrid reliably captures perturbation-induced changes in *m/z* spectra, we next examined effects on the annotated lipid composition. Importantly, although our analysis strategy ensured that most *m/z* values likely correspond to bona fide cellular biomolecules (Figure 1E), only a subset could be annotated to a known lipid species (Figure 2D), reflecting a well-recognized limitation of mass spectrometry-based lipidomics^25^. Using an expert-curated reference list (Table S3), 182 of the 1,377 filtered *m/z* values (13.2%) matched a known ionized lipid species within a 10-ppm mass tolerance (Methods). For some, multiple ionization states collapse on the same molecule (Figure 2D). In total, we could thus annotate 158 unique lipid species, spanning 18 different lipid subclasses. Notably, some of these subclasses (e.g., DGs, PAs and MGs) might also stem from laser-induced fragments of larger lipids^26^. Accordingly, while informative, changes to these annotated species might reflect alterations in the abundance of their precursor lipids.

Using these annotated lipid species, we next assessed changes in lipid composition due to the gene knockouts (log_2_FC knockout / intergenic controls) (Table S4). Different knockouts affected the lipid subclass abundancies in distinct ways (Figure 2E). As expected based on their biosynthetic relationship, ceramides (Cer) and their glycosylated derivatives, dihexosylceramide (Hex2Cer) showed similar perturbation profiles^27^. Likewise, changes to phosphatidylcholine (PC) and sphingomyelin (SM) abundancies were highly concordant, suggesting that their homeostasis is under control of shared regulatory components or pathways. In contrast, apparently related lipid subclasses such as lysophosphatidyl-inositols (LPI) and lysophosphatidyl-ethanolamines (LPE) showed more distinct perturbation profiles, suggesting divergent regulatory mechanisms. While this aggregated analysis reveals overall trends, individual species often also exhibit specific responses to gene knockouts.

We therefore evaluated how each individual lipid species responded to the 143 gene knockouts (Figure 2F). While closely related species often behaved similarly, we also observed clear differential effects between species within the same subclass. The phosphatidylethanolamines PE 36:5 and PE 40:4 were for example anti-correlated (r = -0.21), suggesting that several knockouts have opposing effects on their abundance. While a few knockouts such as ELOVL5 and PLD2 affected only a limited subset of lipid species, most perturbations significantly altered the abundance of multiple species. Some gene knockouts show a global effect on lipid abundance, either positively (e.g. GPAM or PNPLA2) or negatively (e.g. ACOT4 or PTDSS1), while others (e.g. LPCAT3 or ACOX1) significantly reshape the lipidome in both directions.

To assess whether LipoGrid, which uses MALDI-MSI to measure the lipid profile of single cells, reliably captures the impact of gene perturbations on the lipidome, we carried out an orthogonal validation. We knocked down eight target genes using siRNA and profiled the lipidome of each knockdown and of matched controls by quantitative bulk lipidomics using LC-MS^28^ (Methods). Reassuringly, the perturbation effects, quantified as the log_2_FC of 57 lipid species that changed significantly (P < 0.05), correlate significantly between the LipoGrid and bulk measurements (r = 0.52, P = 3.6 × 10⁻⁵) (Figure S5). Together, these results demonstrate that LipoGrid reliably captures perturbation-induced changes in the cellular lipidome at the level of individual lipid species.

### Recapitulating substrate-specific lipid remodeling

To evaluate whether LipoGrid faithfully captures known lipid-modifying activities, we focused on target genes that encode enzymes with a well-established substrate and for which therefore the expected impact on specific lipid species is clearly defined. First, to probe substrate specificity within the long-chain acyl-CoA synthetase (ACSL) family, we compared ACSL3 and ACSL4 knockouts across diacyl glycerophospholipids and DG, stratifying species by acyl-chain saturation^29^. Saturated and monounsaturated fatty acid (sat/MUFA)-containing species were defined as diacyl glycerolipids carrying up to one double bond, whereas polyunsaturated fatty acid (PUFA)-containing species were diacyl species with at least four double bonds. With this stratification, ACSL3 knockout more strongly depleted sat/MUFA-than PUFA-containing phospholipids (median log_2_FC -0.16 vs -0.08, respectively, P = 0.032) while ACSL4 knockout displayed the opposite pattern, more strongly depleting PUFA-than sat/MUFA-containing phospholipid (median log_2_FC -0.46 vs -0.25, respectively, P = 0.0033, Figure 3A). This pattern mirrors the known substrate preferences of the two enzymes: ACSL4 preferentially activates PUFAs for esterification into phospholipids, whereas ACSL3 favors saturated and MUFA species^29,30^.

**Figure 3.**
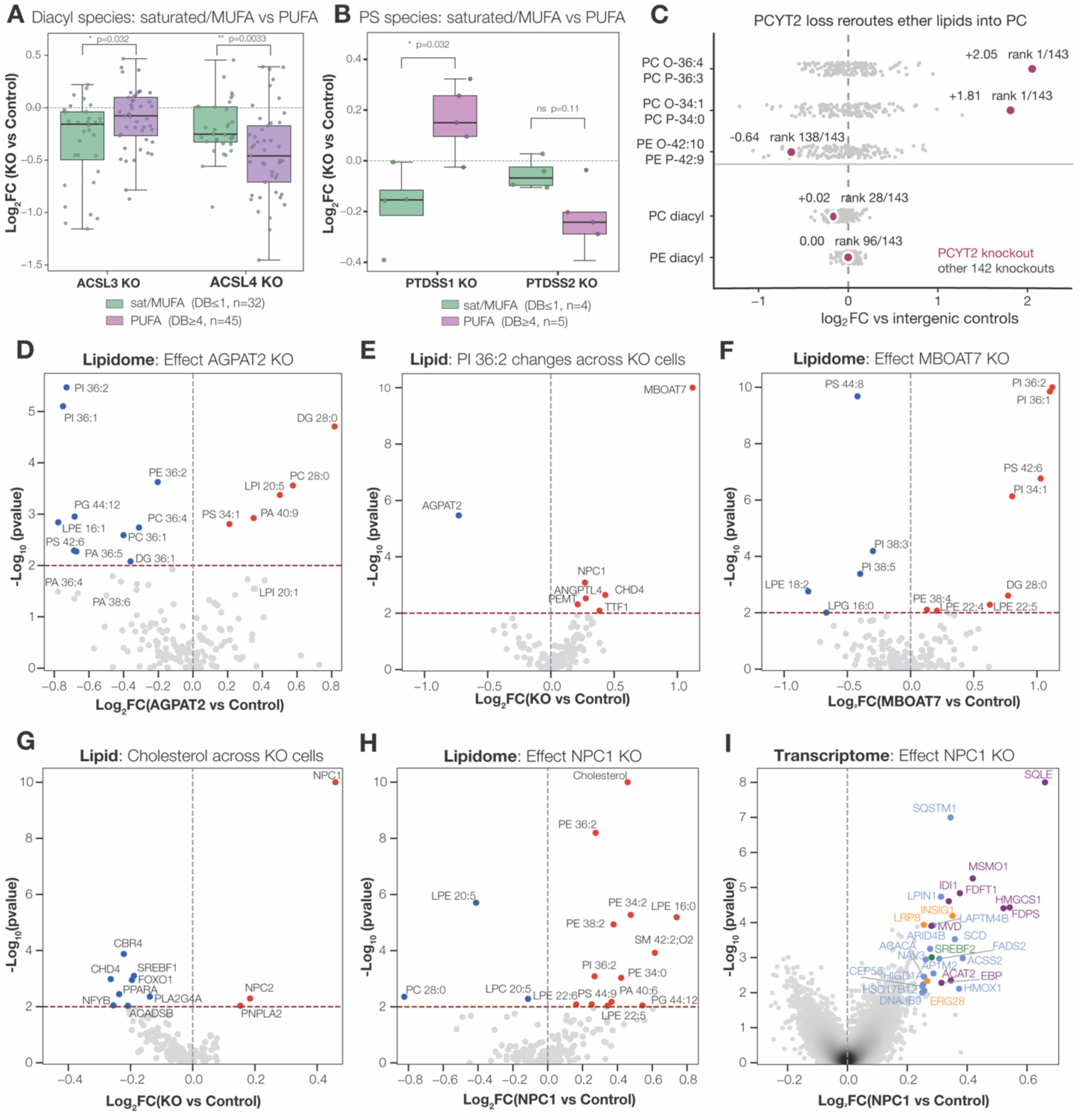
LipoGrid recapitulates substrate preferences and molecular disease phenotypes. (A-B) Boxplots visualizing the log_2_ fold change in mean intensity for selected substrates following specific gene knockouts relative to intergenic controls. (A) Effect of ACSL3 and ACSL4 knockout on all measured saturated/MUFA (DB ≤ 1, n=32) versus PUFA (DB ≥ 4, n=45) diacyl lipid species. (B) Effect of PTDSS1 and PTDSS2 knockout on all measured saturated/MUFA (DB ≤ 1, n=4) versus PUFA (DB ≥ 4, n=5) PS lipid species. (C) PCYT2 loss reroutes ether lipids into PC. Log₂ fold change in mean lipid intensity following gene knockout relative to intergenic controls. Each grey dot is one of the 143 target genes; PCYT2 is highlighted in dark pink with its value and rank. Bottom two rows show the median across all diacyl PC (n = 24) and PE (n = 20) species as a specificity control. (D-H) Volcano plots showing the log_2_ fold change in lipid abundance for each knockout relative to intergenic controls (x-axis) and the corresponding statistical significance, expressed as in -log10 (P value), based on a two-sided Mann–Whitney U test (y-axis). (D) Effect of AGPAT2 knockout on the lipidome. (E) Impact of every knockout on PI 36:2 levels. (F) Effect of MBOAT7 knockout on the lipidome. (G) Impact of every knockout on Cholesterol levels. (H) Effect of NPC1 knockout on the lipidome. (I) Volcano plot visualizing the effect of NPC1 knockout on the transcriptome. Upregulated genes involved in cholesterol biosynthesis are colored; enzymes in purple, regulators in orange and the key transcription factor in green.

A second case where substrate channeling predicts species-level selectivity are the phosphatidylserine synthases PTDSS1 and PTDSS2. Both enzymes catalyze head-group exchange to generate phosphatidylserine (PS), but draw from distinct phospholipid donors and are hence expected to feed PS pools with distinct acyl signatures^31,32^. To test this using a consistent analytical framework, we applied the same strict stratification threshold utilized in our ACSL analysis, defining saturated/MUFA-rich species as those containing ≤1 double bond and PUFA species as those with ≥4 double bonds, which captured 9 of our measured PS species. Consistent with PTDSS1 channeling saturated fatty acids/MUFA-rich acyl profiles into PS, its knockout selectively depleted saturated/MUFA-PS while PUFA-PS accumulated (P = 0.032, Figure 3B). PTDSS2 knockout instead showed a trend toward PUFA-PS depletion (median log_2_FC -0.24), in line with PTDSS2 generating PUFA-rich PS^31,32^. Across all 143 perturbations, PTDSS2 knockout produced the largest PS depletion in the screen (median log₂FC -0.20 across the 12 measured PS species), and depleted PS more than the remainder of its own lipidome (P = 0.011). PTDSS1 loss, by contrast, left the total PS pool unchanged (rank 89 of 143), consistent with the two enzymes supplying PS pools of different size and acyl composition.

A third case revealed rerouting between head-group branches. PCYT2 catalyzes the rate-limiting step of the CDP-ethanolamine branch of the Kennedy pathway, which supplies both diacyl- and ether-linked PE^33^. Its knockout left the bulk diacyl PE and PC pools unchanged (median log₂FC 0.00 and +0.02) but produced the two largest lipid inductions of any perturbation in the screen: both detected ether-PC species increased more than threefold and ranked first of all 143 knockouts (log₂FC +2.05 and +1.81; P = 2.8 × 10⁻^20^ and 2.1 × 10⁻^20^) while the single ether-PE species decreased (Figure 3C). This reciprocal shift is consistent with the alkyl/alkenyl-glycerol backbone being diverted into the CDP-choline branch when the ethanolamine branch is blocked, with the diacyl PE pool spared by continued synthesis from phosphatidylserine decarboxylation^32,34^.

Together these three cases show the screen recovering enzyme specificity at the level on which each enzyme acts, acyl-chain saturation for ACSL3 and ACSL4, head-group donor for PTDSS1 and PTDSS2, and branch selection for PCYT2, indicating that LipoGrid resolves substrate preference rather than only the direction of a perturbation’s effect.

### Resolving gene-lipid relationships at lipid species resolution

We further evaluated whether LipoGrid faithfully captures functional relationships between genes and lipids by checking how the knockout of genes encoding proteins with established roles in lipid metabolism affects the entire measured lipidome. Similar as before, we quantified lipid abundance changes as log₂ fold changes relative to intergenic controls and assessed significance using a nonparametric Mann–Whitney U test (Methods, Table S4). We first examined *1-acylglycerol-3-phosphate O-acyltransferase 2* (AGPAT2), the gene mutated in congenital generalized lipodystrophy type 1 (CGL1; Berardinelli-Seip syndrome), an autosomal recessive disorder marked by near-complete loss of adipose tissue, severe insulin resistance and hypertriglyceridemia^35^. AGPAT2 catalyzes the acylation of lysophosphatidic acid (LPA) to phosphatidic acid (PA), a committed step in de novo triglyceride and phospholipid synthesis, and its loss is thought to impair the lipid production required for adipocyte storage^35,36^. Its knockout produced this signature in the lipidome: the direct enzymatic product PA was largely depleted, most strongly for polyunsaturated species (e.g. PA 36:4, PA 36:5 and PA 38:6 reduced by 42%, 37% and 29%; P = 0.032, 0.0053 and 0.038), while lysophospholipids upstream of the block accumulated (LPI 20:5 and LPI 20:1 increased by 42% and 28%; P = 4.2 × 10⁻⁴ and 0.029), consistent with a stalled acylation step (Figure 3D). This loss of PA propagated to its downstream phospholipids: an earlier study showed that AGPAT2 overexpression increases PI 36:1 and 36:2 abundance^37^, and correspondingly, our data showed that AGPAT2 knockout significantly reduced both species by 41% and 40%, respectively (P = 7.9 × 10⁻⁶ and 3.4 × 10⁻⁶; Figure 3D). Across all perturbations, AGPAT2 knockout produced by far the largest reduction in PI 36:2 of any screened gene (−40% versus ≤ 27% for the next-strongest perturbations), indicating a predominant role in PI 36:2 biosynthesis in our screen (Figure 3E). In contrast, several knockouts increased PI 36:2 levels, with MBOAT7 loss showing the strongest effect. MBOAT7 encodes membrane-bound O-acyltransferase 7 (LPIAT1), a lysophospholipid acyltransferase that selectively re-acylates lyso-PI with arachidonoyl-CoA and thereby channels the PI pool towards polyunsaturated species^38^. Its loss is therefore predicted to remodel PI directionally, depleting polyunsaturated PI while allowing saturated and monounsaturated species that escape re-acylation to accumulate^39^. LipoGrid reproduced precisely this reciprocal signature: MBOAT7 knockout increased PI 36:2 more than twofold and similarly raised PI 36:1 and PI 34:1 (log₂FC = 1.12, 1.1 and 0.7; P = 1.7 × 10^⁻15^, 1.4 × 10^⁻10^ and 7.4 × 10⁻^7^), while significantly decreasing the polyunsaturated species PI 38:3 and PI 38:5 (log₂FC = -0.3 and -0.4; P = 6.5 × 10⁻5 and 4.2 × 10^-4^) (Figure 3F).

Notably, AGPAT2 and MBOAT7 shape the same PI species through distinct routes. AGPAT2 acts upstream, supplying the phosphatidic acid precursor from which PI is synthesized de novo, whereas MBOAT7 acts downstream, exchanging acyl chains within the mature PI pool. Their opposing effects on PI 36:1 and PI 36:2 therefore reflect the two arms of PI homeostasis, synthesis and remodeling, rather than competing activities on a shared step, illustrating how LipoGrid can separate biosynthetic from remodeling control of an individual lipid species.

### Recapitulating the NPC1 loss-of-function phenotype

Beyond a gene-centered analysis, LipoGrid data also reveals genetic regulators of specific lipid species. As an example, we focused on cholesterol, an essential membrane lipid dysregulated in diverse cardiovascular, metabolic and neurodegenerative diseases^40^. Perturbations of 11 genes significantly altered cholesterol abundance, but none by more than 38%, suggesting pleiotropic control of cholesterol levels. Eight knockouts significantly lowered cholesterol levels (Fig. 3G), including genes encoding transcriptional regulators (*CHD4, NFYB, PPARA, FOXO1* and *SREBF1*) and ancillary lipogenic enzymes (*CBR4, ACADSB* and *PLA2G4A*). This suggests that a reduced activity of the overall lipogenic transcriptional program lowers free cholesterol. Conversely, knockout of *NPC1, NPC2* and *PNPLA2* increased cholesterol abundance. While PNPLA2 regulates triglyceride lipolysis and lipid droplet homeostasis, which could as such influence cholesterol storage and mobilization^41^, NPC1 and NPC2 directly mediate intracellular cholesterol trafficking^42^. Loss-of-function mutations in either gene cause Niemann–Pick disease type C, which is characterized by lysosomal accumulation of cholesterol and other lipids^43,44^. Among all perturbations, NPC1 loss produced the strongest cholesterol increase (37.5% more cholesterol, P = 2 × 10⁻^15^) and caused accumulation of additional lipid species (Figure 3H), consistent with the established Niemann-Pick disease type C phenotype^43^.

Earlier work has linked cholesterol accumulation in NPC-deficient cells to impaired export of unesterified cholesterol from late endosomes and lysosomes^45^. Although cholesterol accumulates in lysosomes, its reduced delivery to the endoplasmic reticulum activates the SREBP2-mediated cholesterol homeostatic response^46^. As we also measured the transcriptional responses for each knockout (CROP-seq) in parallel with the lipidome changes (Figure S6, Methods), we next asked if NPC loss elicited this canonical feedback loop. In the scRNA data, *NPC1* knockout significantly altered expression of only 28 genes (|log_2_FC| > 0.25, P < 0.01) (Fig. 3I). Notably, 13 upregulated genes encoded members of the cholesterol biosynthesis pathway (GO REAC, P = 4.53 × 10^-19^). These included the transcription factor *SREBF2*, the master regulator of cholesterol homeostasis, together with multiple enzymes in the cholesterol biosynthetic pathway, including *SQLE, HMGCS1, FDPS, MSMO1, FDFT1, ACAT2, IDI1, EBP,* and *MVD*. Overall, the lipidomic and transcriptomic data demonstrate that LipoGrid faithfully recapitulates the NPC loss-of-function phenotype.

### Mapping transcriptional responses to lipid-associated gene perturbations

As the NPC1 example illustrates, lipid level changes induced by genetic perturbations may reflect both direct metabolic effects and indirect, compensatory changes. The latter may arise when cells sense altered lipid abundance and accordingly adjust the expression of biosynthetic or catabolic enzymes. To more systematically assess these transcriptional responses, we performed batch Gene Set Enrichment Analysis (GSEA) using a curated compendium of 824 lipid-associated gene sets compiled from multiple databases (Methods). We identified 91 discrete gene sets that were enriched in at least one knockout (Table S5, Figure S7). Interestingly, while some knockouts, such as *FADS2* and *DGAT1*, produced few or no significantly enriched terms, most induced overt transcriptional changes in lipid-related transcriptional programs (Figures 4A and S7). Some responses were highly specific. For example, knockout of *B4GALT6*, which encodes β-1,4-galactosyltransferase 6, selectively induced expression of genes involved in glycosphingolipid binding. Knockout of *APOE*, encoding a component of the lipid transport system, upregulated genes associated with isoprenoid binding and downregulated those associated with glycolipid binding.

**Figure 4.**
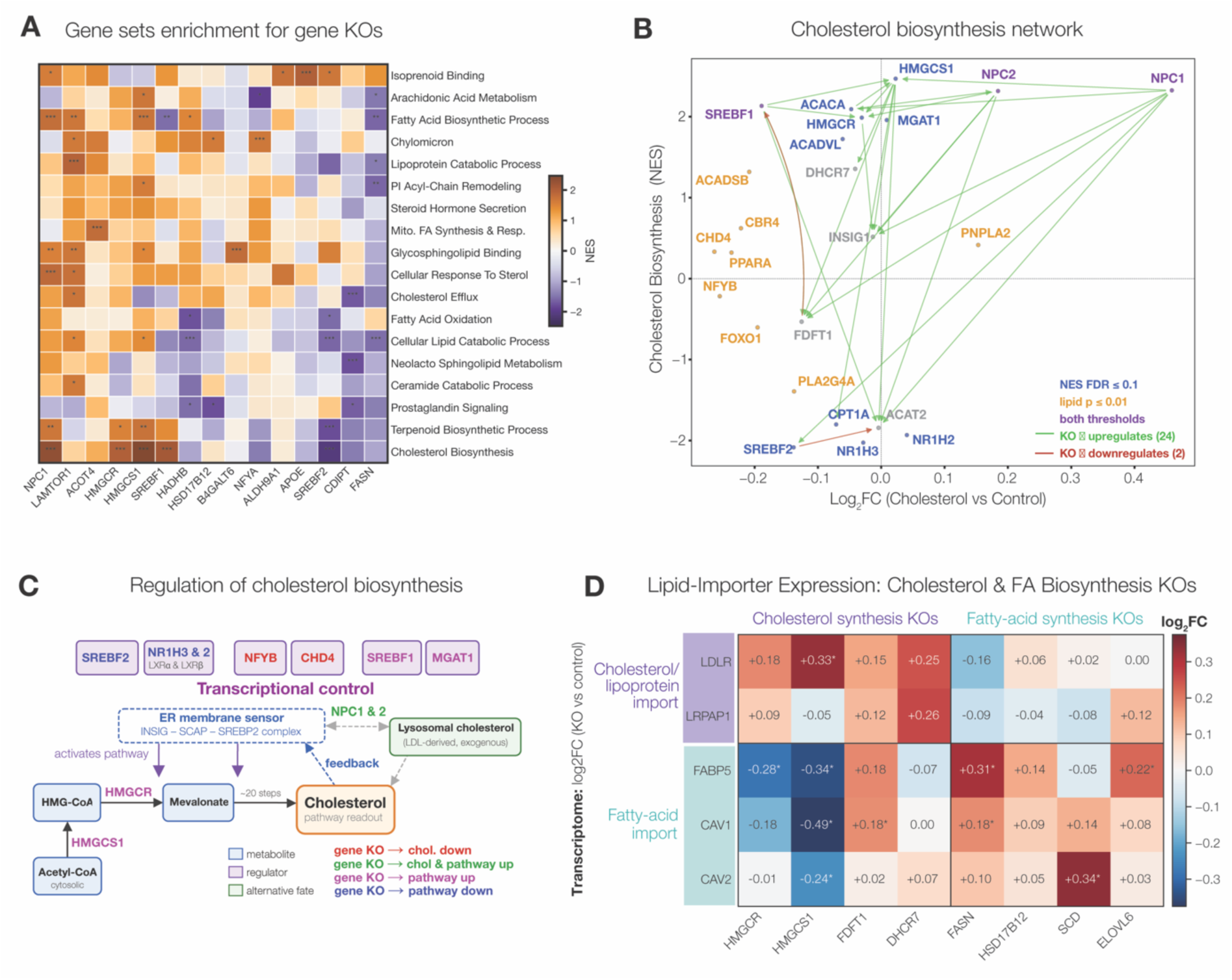
Compensatory transcriptional responses to lipid metabolic gene knockouts. (A) Heatmap showing the normalized enrichment score (NES) for a subset of the lipid related gene sets across the indicated gene knockouts, full heatmap in Figure S7. (B) Connected dot plot showing the relationship between the log_2_ fold change in cholesterol abundance following gene knockout relative to intergenic controls (x-axis) and the normalized enrichment score (NES) of the cholesterol biosynthesis gene set (y-axis). Blue gene names indicate knockouts that significantly altered cholesterol biosynthesis gene expression (NES FDR < 0.1), yellow indicate knockouts that significantly affected cholesterol abundance (P < 0.01), and purple if they pass both thresholds, grey represent other cholesterol-related genes. Arrows connect genes whose knockout significantly (P < 0.01) altered the transcription of the indicated target genes. (C) Metabolic pathway diagram illustrating part of the cholesterol biosynthesis pathway. Key regulators included in the screen are colored according to the effects of their knockout: red, reduced cholesterol levels; green, increased cholesterol levels and biosynthesis gene expression; purple, induction of cholesterol biosynthesis genes; and blue, reduced expression of cholesterol biosynthesis genes. (D) Heatmap showing the log_2_ fold change in expression of genes involved in extracellular cholesterol and fatty acid uptake following knockout of cholesterol and fatty acid biosynthesis enzymes, relative to intergenic controls.

Other knockouts produced a broader remodeling of lipid-related transcriptional programs. This was particularly evident for transcription factors. For example, knockout of *SREBF2*, which encodes the master regulator of cholesterol homeostasis^46^, reduced expression of genes associated with terpenoid biosynthesis, cholesterol metabolism, and lipid catabolism (Figure 4A). Similarly, knockout of the more general transcription factor NFYA increased expression of genes linked to chylomicron production while reducing expression of genes involved in arachidonic acid metabolism, steroid dehydrogenase activity, and cholesterol metabolism (Figure S7). Broader transcriptional responses were not limited to transcription factors. Knockout of metabolic enzymes such as ALDH9A1, HMGCS1, HSD17B12, and HADHB also induced substantial transcriptional changes, likely reflecting secondary consequences of disrupted metabolic fluxes or altered lipid homeostasis. Overall, most knockouts significantly altered the expression of genes associated with defined metabolic processes. These results demonstrate that LipoGrid captures coordinated transcriptional responses to lipid perturbations and reveals widespread remodeling of lipid-associated transcriptional programs.

### Lipid metabolic perturbations trigger compensatory feedback responses

Interpreting a perturbation’s impact on the lipidome requires accounting for accompanying transcriptional responses. For example, knockout of key cholesterol biosynthesis enzymes such as the mevalonate-pathway enzymes HMGCS1 and HMGCR had surprisingly little effect on the overall cholesterol abundance. However, batch GSEA revealed that both knockouts strongly induced expression of cholesterol biosynthesis genes (NES +2.47 and +1.99), consistent with compensatory transcriptional feedback that buffers loss of these enzymes (Figure 4A).

To examine cholesterol homeostasis across perturbations, we compared changes in cholesterol abundance with the transcriptional activity of the cholesterol biosynthesis program (Figure 4B). Several knockouts strongly induced expression of this gene set. In addition to HMGCS1 and HMGCR, also knockout of the cholesterol trafficking genes *NPC1* and *NPC2* (NES = +2.33 and +2.32) and central controllers of lipid metabolism such as *SREBF1* and *ACACA* (NES = +2.13 and +2.09) induced expression of the cholesterol biosynthesis program. Conversely, the same program was downregulated by knockout of the master sterol transcription factors *SREBF2*, *NR1H3* (encoding LXRα) and *NR1H2* (encoding LXRβ) (NES = -2.08, -2.03 and -1.93), identifying this transcription factor backbone (SREBF2 + LXRα/β) as driving cholesterol-responsive transcription^47^. Additionally, our screen reveals which knockouts drive the up- or downregulation of other genes within the screen. For the cholesterol biosynthesis pathway, we observed a striking predominance of positive effects (24 upregulated versus 2 downregulated connections; Figure 4B), whereby knockout of a single enzyme led to significant induction of multiple other pathway components. Together, these patterns match the canonical SREBP2 feedback model. Knockout of biosynthetic enzymes, or of the NPC1/NPC2 route that delivers lysosomal cholesterol to the ER, depletes the ER sterol pool. This triggers the release of SCAP-SREBP2 from INSIG and its transport to the Golgi, where SREBP2 cleavage and transcriptional activation induces the cholesterol biosynthesis program^46^ (Figure 4C). Thus, LipoGrid captures the feedback logic of cholesterol homeostasis and reconstructs key regulatory relationships within the pathway in a single experiment.

### Lipid synthesis inhibition triggers compensatory lipid import

Beyond the transcriptional upregulation of alternative enzymes within a pathway to compensate for the loss of a single catalytic step, cells can also restore lipid flux by increasing the import of specific lipid subclasses from the extracellular environment. A better understanding of these compensatory routes is crucial, as exogenous lipid scavenging is a well-documented escape mechanism by which tumors circumvent pharmacological blockade of de novo fatty acid synthesis, for instance upon inhibition of fatty acid synthase (FASN) or acetyl-CoA carboxylase (ACC)^48,49^.

To investigate this, we examined how the expression of lipid-import genes responded to loss of key enzymes in the cholesterol and fatty-acid synthesis pathways (Methods). Only a small number of import genes were sufficiently expressed in A549 cells, with the *low-density lipoprotein receptor (LDLR)* and *LDL-receptor-related protein-associated protein 1 (LRPAP1)* on the cholesterol/lipoprotein side and *fatty-acid-binding protein 5 (FABP5)*, *caveolin-1 and 2 (CAV1 and CAV2)* on the fatty-acid side (Figure 4D). The clearest signal was the specific upregulation of *LDLR* upon knockout of the cholesterol biosynthesis enzymes. LDLR is a cell-surface receptor that mediates uptake of cholesterol-rich lipoproteins from the environment^50^, and its specific upregulation indicates that cells try to compensate the loss of cholesterol synthesis by exogenous import. HMGCS1 loss reproduced the cholesterol-deprivation program cleanly, by not only inducing *LDLR*, but also lowering the fatty acid import genes. On the fatty-acid axis, *FABP5*, a small cytosolic chaperone that binds long-chain fatty acids^51^, was upregulated following perturbation of genes involved in fatty-acid metabolism, with the strongest increases observed upon knockout of *FASN* (log_2_FC +0.31, P = 0.015) and *ELOVL6* (log_2_FC +0.22, P = 0.019) (Figure 4D). In contrast, knockout of the desaturase *SCD* did not induce *FABP5* expression but instead increased *CAV1* (log_2_FC +0.14, P = 0.22) and *CAV2* (log_2_FC +0.34, P = 3.8 × 10⁻⁴) expression, which encode the principal structural components of caveolae^52^. Caveolins organize lipid microdomains and facilitate membrane trafficking, lipid uptake, lipid droplet formation and neutral lipid storage. *CAV1* was targeted in our screen, and its knockout similarly depleted the neutral glycerolipid pool, with DGs and MGs showing the strongest reductions (median log2FC -0.38 and -0.37, respectively), compared with a more modest decrease in PCs (−0.23) and minimal effects on other phospholipids. This pattern is consistent with the established role of caveolin-1 in lipid droplet formation and neutral lipid import and storage^52,53^.

Overall, these data demonstrate that disruption of lipid biosynthetic enzymes induces a compensatory upregulation of genes involved in lipid uptake. Notably, this response is pathway-specific, as perturbation of cholesterol biosynthesis selectively increased the expression of cholesterol uptake genes, whereas disruption of fatty-acid biosynthesis preferentially induced fatty-acid uptake and trafficking genes (Figure 4D). This distinction highlights the existence of coordinated, lipid class-specific homeostatic programs that couple impaired biosynthesis to enhanced extracellular lipid acquisition.

By integrating lipidomic and transcriptomic responses to perturbation, LipoGrid reveals not only the regulatory mechanisms of lipid metabolism but also the compensatory programs that act to restore homeostasis, establishing a generalizable approach for connecting genotype to lipid phenotype in health and disease.

## Discussion

In this study, we combined a pooled CRISPR knockout screen with grid-assisted single-cell mass spectrometry and spatial transcriptomics to generate a unique, internally controlled multi-omic perturbation dataset linking targeted disruption of lipid metabolic genes to their downstream lipidomic and transcriptional consequences. To our knowledge, this is the first study to profile the perturbed cellular lipidome and transcriptome at single-cell resolution across a targeted set of 143 lipid metabolism genes. To facilitate exploration of this resource, we provide an interactive graphical user interface that enables intuitive querying and visualization of gene-, lipid-, and transcript-level perturbation effects.

LipoGrid recapitulated several established lipid regulatory relationships, demonstrating its ability to resolve gene-lipid associations at molecular resolution. ACSL4 knockout reproduced its known substrate selectivity, preferentially remodeling PUFA-containing over MUFA-containing phospholipids. Similarly, disruption of the NPC1/NPC2 cholesterol transport pathway led to the expected accumulation of cholesterol, while AGPAT2 and MBOAT7 loss converged on the same PI 36-series species from opposite ends of the pathway: AGPAT2 by restricting the phosphatidic acid supply for de novo PI synthesis, and MBOAT7 by diverting lyso-PI away from arachidonoyl re-acylation, shifting the PI pool from polyunsaturated towards monounsaturated species. Notably, NPC1 and AGPAT2 perturbations also recapitulated the characteristic molecular signatures associated with their respective loss-of-function disease states, further supporting the biological fidelity of LipoGrid-derived lipidomic profiles.

By combining lipidomic and transcriptomic readouts in single cells, LipoGrid provides mechanistic insight inaccessible to either modality alone. A recurring theme in our data is the robustness of cellular lipid metabolism to genetic perturbation, buffered by compensatory transcriptional and metabolic rewiring. This is exemplified by the coordinated induction of cholesterol biosynthesis genes following knockout of multiple pathway enzymes, a response that likely acts to restore cholesterol homeostasis. Beyond transcriptional compensation within biosynthetic pathways, cells also adapted to reduced lipid flux by increasing uptake of extracellular lipids. Perturbation of cholesterol or fatty acid biosynthesis enzymes induced expression of genes involved in the uptake of extracellular cholesterol or fatty acids, respectively. In the context of lipid-rich 10% FBS culture conditions, this enhanced uptake likely mitigated defects in de novo lipid synthesis, contributing to the modest phenotypic effects observed for some perturbations, such as the limited impact of FASN knockout on the free fatty acid pool. Together, these findings reveal that single-gene perturbations are buffered by interconnected uptake, recycling, and remodeling pathways, highlighting that metabolic phenotypes are shaped not only by the targeted enzyme but also by the broader regulatory network in which it operates.

Understanding these adaptive mechanisms has important therapeutic implications. In oncology, lipid metabolic dependencies are increasingly being explored as vulnerabilities, and identifying the compensatory pathways that enable tumor cells to evade metabolic inhibition will be critical for designing rational combination therapies^48,54^. Similar principles apply to neurodegenerative disorders, where disrupted lipid trafficking and membrane lipid homeostasis are emerging contributors to disease progression^55^.

As with any emerging platform, this study highlights opportunities for further development. Because CRISPR knockout penetrance varies across cells, the observed effect sizes are likely conservative and should improve with higher editing efficiencies. Although our current panel of 153 genes captures key aspects of lipid biology, LipoGrid is readily scalable toward the ∼1,200 genes implicated in mammalian lipid metabolism, enabling systematic mapping of regulatory and compensatory networks across diverse cell types and physiological contexts. Such datasets could support predictive models of lipid remodeling in response to genetic perturbation, providing a foundation for applications ranging from molecular diagnosis of inherited lipid disorders and lipidomic stratification of cancer to the rational development of metabolic therapies. Comprehensive maps of compensatory regulatory networks may also guide combination strategies to overcome metabolic adaptation and therapeutic resistance.

In conclusion, LipoGrid establishes a scalable framework for dissecting how gene perturbations propagate through metabolic networks at single-cell resolution. By systematically capturing both lipidomic outputs and the compensatory transcriptional programs that buffer them, the approach offers a path toward a mechanistic understanding of lipid-metabolic plasticity and, ultimately, toward identifying the network-level vulnerabilities that can be exploited therapeutically in cancer, neurodegeneration, and other diseases of lipid dysregulation.

## Acknowledgements

J.J. and A.S. are funded by the Leuven Future Fund (LISCO-BIOMED), Opening the Future Fund, KU Leuven ID-N (3E210655 COLUMBO). P.V.M. and E.D.R. are funded by a PhD fellowship from Fonds Wetenschappelijk Onderzoek – Vlaanderen (FWO(11K6222N) and (1SHD124N) respectively). N.R. by KU Leuven C1 grants C14/21/095 (InterAction) and C14/25/154 (GRAN’PA). This work was supported by an FWO-SBO grant LIPOMACS (S001623N) to J.V.S., a KU Leuven Core Facility Translational Grant. L.V., A.S. and J.V.S. are supported by HORIZON-MSCA-2022-DN-01, project number 101120283 -PROSTAMET. T.V., K.Va. and N.V. are supported by KU Leuven (C14/22/125, Leuven Future Fund), the Research Foundation Flanders (FWO, G005923N and I009724N) and VLIR (Vlaamse Veerkracht, PRISMO). K.Ve and J.L. are funded by KUL C2 (C24E/20/035) and KUL iBOF (iBOF/23/005).

## Author contributions

B.T., A.S. and J.J. conceptualized the project. P.V.M. prepared CROP-seq library, performed cell culture, scRNA-seq, and optimized the wet lab procedures under supervision of B.T., N.R. optimized and performed MALDI-MSI under supervision of J.V.S., E.D.R. optimized and generated the micropatterned grids under supervision of K.Ve., J.B. and J.L., N.V. optimized and performed 10X Xenium under supervision of K.Va and T.V., L.V. designed the preprocessing pipeline for MALDI-MSI and alignment. D.W. performed segmentation on Xenium. J.J. performed the data analysis. J.V.S. and N.R. provided expertise in lipid metabolism and interpretation of lipid metabolic perturbations. J.J. wrote the paper together with P.V.M, B.T., A.S and with inputs from all authors. J.J. supervised the project with B.T. and A.S.

## Declaration of generative AI and AI-assisted technologies in the manuscript preparation process

During the preparation of this work, the author(s) used Claude (Opus 4.8) and ChatGPT for assistance with coding and refining small parts of the text. The author(s) reviewed and edited the output as needed and take full responsibility for the content of the published article.

## Material and methods

### Data availability

Raw and processed 10x Xenium and sc-RNA-seq (CROP-seq) data are available on GEO using the following accession number: GSE339212.

Raw and processed MALDI-MSI data, together with the coordinate file that links spatial transcriptomics with spatial lipidomics are available on zenodo: (https://doi.org/10.5281/zenodo.21456908).

Code and notebooks are available on https://github.com/sifrimlab/LipoGrid.

### Cell culture

A549 cells (ATCC) were maintained in DMEM (ThermoFisher, 41965062) supplemented with 10% FBS (ThermoFisher, A5256701) and 5% penicillin–streptomycin (ThermoFisher, 15140122) at 37 °C in 5% CO₂. HEK293T cells (ATCC) were maintained in IMDM (ThermoFisher, 12440053) supplemented with 10% FBS and 5% penicillin–streptomycin at 37 °C in 5% CO₂. Cells were passaged using 0.25% Trypsin–EDTA (ThermoFisher, 25200056). Mycoplasma testing was performed monthly using the MycoAlert® Mycoplasma Detection Kit (Lonza, LT07-318).

### Generation of doxycycline-inducible Cas9 monoclonal A549 cells

Cells were transduced with a lentivirus encoding the Edit-R Inducible Lentiviral Cas9 Nuclease (Horizon Discovery, CAS11229) and selected with 20 μg/mL blasticidin (InvivoGen, 38220000) for 14 days. Single-cell clones were isolated by sorting into 96-well plates using a BD FACS Aria Fusion and expanded in conditioned medium before transfer to 6-well plates.

Cas9 expression was validated by immunofluorescence. Cells were permeabilized with 0.2% Triton X-100 in DPBS (ThermoFisher, 14190-094) for 10 min and blocked with 10% normal goat serum (Life Technologies, 50197Z) for 40 min at room temperature. Cells were incubated with mouse anti-Cas9 antibody (Novus Biologicals/Bio-Techne, NBP2-36440SS; 1:500 in 2% BSA/DPBS) for 1 h at 37 °C, followed by Alexa Fluor 488-conjugated goat anti-mouse secondary antibody (Life Technologies, A-11017; 1:500) for 40 min at 37 °C in the dark. Nuclei were counterstained with 300 nM DAPI in DPBS for 5 min. Images were acquired using a Leica TCS SP5 confocal microscope.

### Pooled CRISPR gRNA library construction and cell transduction

For each gene, three gRNAs were designed based on VBC score and compatibility with the 10x Xenium spatial transcriptomics platform (preferred junction and off-target criteria). Oligonucleotides containing the gRNA sequences and homology arms for the CROP-seq Guide Puro vector (Addgene #86708) were synthesized as an oPool (Integrated DNA Technologies).

Library cloning was performed following the CROP-seq protocol described by Datlinger *et al*^11^. Briefly, the CROP-seq vector was digested with BsmBI (Bioke, R0739S) and the backbone was purified using the NucleoSpin Gel & PCR Clean-up kit (Macherey-Nagel, 740609.50). The pooled gRNA library was assembled into the backbone using NEBuilder HiFi DNA Assembly Master Mix (NEB, E2621S) and transformed into Endura electrocompetent cells (Lucigen, 60242). Plasmid DNA was isolated using the NucleoBond Xtra Maxi EF kit (Fisher Scientific, 11982482).

Lentivirus was produced by transfecting HEK293T cells with the gRNA plasmid pool together with pMDLg/pRRE, pRSV-Rev, and pMD2.G packaging plasmids (Addgene #12251, #12253, #12259) using Lipofectamine 3000 (ThermoFisher, L3000015). Viral titers were determined by puromycin (ThermoFisher, A1113803) selection and quantification of resistant colonies following Crystal Violet staining (Sigma-Aldrich, 61135).

A549-iCas9 cells were transduced with the lentiviral gRNA library at a multiplicity of infection of 0.2. Positively transduced cells were selected with 60 μg/mL puromycin. Cas9 expression was induced during selection by addition of 5 μg/mL doxycycline (VWR, J60579.14). After 9 days, cells were prepared for seeding on the Xenium and microscopic slide.

### Fabrication of PDMS micropillar stamp for microcontact printing

3-inch CZ silicon wafers were purchased from Microchemicals and SU-8 2015 from Chimie. Polydimethylsiloxane (PDMS) was prepared using a Sylgard 184 silicone elastomer kit (Dow Corning, supplied by Farnell).

The micropillar stamp design, used to create the grid of fibronectin patches, was created using AutoCAD, KLayout and CleWin software. The micropillars were designed as a grid of 30 µm x 30 µm square features with a height of 20 µm and 40 µm spacing in between, and the overall grid size matched the Xenium working area (22.45 x 10.45 mm), yielding a total of 47878 patches within the addressable area. An SU-8 master mold for the micropillar stamp was fabricated using standard soft-photolithography. Briefly, a 3-inch silicon wafer was rinsed with acetone and isopropyl alcohol, followed by air drying and a dehydration bake at 180 °C for 5 min. SU-8 2015 was spin-coated at 500 rpm for 10 s, followed by 2200 rpm for 30 s, to obtain a thickness of 20 µm. The coated wafer was soft-baked at 95 °C for 3.5 min, exposed at 145 mJ/cm2, post-baked at 95 °C for 4.5 min, and finally developed in PGMEA for 3.5 min. The PDMS micropillar stamp was fabricated by replica molding from the SU-8 master mold. PDMS elastomer and curing agent were mixed at a 10:1 mass ratio, degassed for 30 min, and poured onto the SU-8 master mold to a thickness of approximately 7 mm. After a second degassing step to ensure proper filling of all micropillar features, the PDMS was cured overnight at 60°C. The cured PDMS was then peeled from the mold and cut to match the size of the Xenium working area. To improve mechanical stability during microcontact printing, an additional ∼5 mm border of unpatterned PDMS was retained around the micropillar array, reducing lateral drift and smearing and ensuring clean pattern transfer. The grid corners were marked to facilitate alignment with the Xenium working area, as contrast between patterned and unpatterned regions was low.

### Microcontact printing of fibronectin grid on Xenium slides

Fluorescein isothiocyanate-labeled fibronectin (fibronectin-FITC; 1 mL, 1 mg/mL) and regular fibronectin (human origin, 1 mg) were purchased from Sigma-Aldrich and stored according to the manufacturer’s instructions. Phosphate-buffered saline (PBS; 1x, pH 7.4) was prepared from powder (Sigma-Aldrich), dissolved in ultrapure water, and sterile-filtered prior to use. Fibronectin-FITC was used for visualization purposes during the optimization phase, while for the final experimental runs, unlabeled fibronectin was used.

Before printing, the PDMS stamp was cleaned using pressurized N2 and scotch tape to remove dust particles. Crucially, the PDMS stamp was not activated with air plasma, as it would prevent transfer of the proteins to the Xenium surface. A volume of 200 µL fibronectin, diluted in PBS to a concentration of 50 µg/mL, was pipetted over the micropillar region. Due to the hydrophobic nature of PDMS and the limited contact area, spreading of the droplet across the micropillar grid using a pipette tip was required to ensure full coverage. The stamp was then incubated for 20 min under low light conditions to allow fibronectin to adsorb. After incubation, the stamp was washed by immersion in deionized water and dried using an N2 pressure gun.

A Xenium slide and its cassette was sterilized by incubation in 70% ethanol followed by 30 min UV exposure. Using the marked corners on the PDMS stamp as a visual aid, the stamp was aligned onto the Xenium working area. One edge was first brought into contact with the slide, after which the stamp was gradually lowered to ensure uniform contact and prevent smearing. A firm single press perpendicular to the surface was applied, after which the stamp was carefully removed. If this step is not followed correctly, the pattern smears out or is stamped multiple times. Protein transfer is instantaneous upon contact, so no incubation time is needed. The patterned Xenium slide was mounted into the cassette from 10x Genomics, and the resulting chamber, covering the working area, was filled with 1 mL PBS to preserve the printed protein features. As such, slides could be stored at 4°C for at least 3 days prior to use.

### Optimization of patch size

We evaluated square patches of 20, 25, 30, 40 and 50 µm with a fixed interpatch spacing of 40 µm to avoid overlap of MALDI-MSI sampling. Smaller patch features (20-25 µm) resulted in unreliable protein transfer, likely due to smearing incomplete stamping. On the other hand, larger patches (40-50 µm) were frequently occupied by multiple cells, drastically reducing the single-cell efficiency of the grid. Hence for A549 cells, patches of 30 by 30 µm with interpatch spacing of 40 µm was optimal. We do note that these parameters are expected to depend on cell size and adhesion properties and should therefore be adapted for other cell types.

### Cell seeding on patterned Xenium slides

Prior to cell seeding, the fibronectin-patterned working area was washed twice with 1x DPBS and once with complete culture medium. Single-cell suspensions were prepared in complete culture medium supplemented with 5 μg/mL doxycycline and filtered through 40 µm Flowmi® cell strainers (Sigma-Aldrich, BAH136800040). Cells were seeded in the working area at concentrations of 102 cells / mm2 (run 1 and 2) and 204 cells / mm2 (run 3 and 4) to achieve 50% and 100% coverage, respectively, where coverage was calculated as ([cells / patches] x 100). To promote uniform adhesion and prevent aggregation, slides were handled gently and movement was minimized following seeding. Cells were incubated overnight to allow adhesion, followed by three gently washes with 750 µL 1× DPBS to remove unbound cells. It should be noted that these parameters are expected to vary with cell adhesion properties and should therefore be optimized for different cell types.

### Preparation slides for MALDI

Patterned cells were fixed with ice-cold 4% paraformaldehyde (Fisher Scientific, 28906) for 15 min at room temperature and washed three additional times with 1× DPBS. They were subsequently washed with an ice-cold solution of 150 mM ammonium formate in water for 10 times 20 seconds and 5 times 10 seconds using a fresh solution each time. Then, the slides were dried under a gentle nitrogen stream for 15 min. The sample was scanned using an FS120 scanner (Braun, Germany). Using the HTX M5 sprayer (HTXImaging, HTX Technologies, LLC, Chapel Hill, US), the matrix N-(1-naphthyl)ethylenediamine dihydrochloride (NEDC) (Sigma-Aldrich, N9125) was sprayed on top of the slides. A 7 mg/ml solution in 70% MeOH was prepared, and 18 layers were sprayed in the CC pattern with 3 mm track spacing at the nozzle temperature of 75°C and velocity of 1200 mm/min, 10 psi nitrogen pressure, and methanol:water (1:1 *v/v*) pump solvent flow of 0.60 ml/min, and plate temperature of 25°C.

### Spatial lipidomics using MALDI-MSI

The MALDI-MSI imaging was performed on a timsTOF fleX MALDI-2 mass spectrometer (Bruker Daltonik, Bremen, Germany).

Slides 1 and 2 were acquired in positive ion mode only. A 26 × 26 µm laser beam scan was used, resulting in a 30 × 30 µm pixel size (step size). MALDI-2 post-ionization was applied to enhance lipid detection. Primary ionization was performed using the Bruker SmartBeam 3D laser (40 shots, 1 kHz, 50% power, single SmartBeam mode with beam scanning). The MALDI-2 laser (NL204–1K-FH, Ekspla) was operated with a 5 µs pulse delay. The *m/z* range was 300–1800. Deflection 1 delta was set to 70 V. Pre-TOF transfer time and pre-pulse storage were 85 µs and 5 µs, respectively.

Slides 3 and 4 were acquired in positive followed by negative ion mode. The laser operated at 200 shots, 10 kHz, and 50% power in single SmartBeam mode with beam scanning. The *m/z* range was extended to 300–2000. Deflection 1 delta was set to –70 V. Pre-TOF transfer time remained 85 µs, with pre-pulse storage increased to 10 µs.

All other instrument parameters were identical between runs: MALDI plate offset 60 V; funnel 1 and 2 RF voltages 300 and 350 Vpp; multipole 400 Vpp; collision cell 1800 Vpp; energy gap 10 eV (quadrupole–collision cell) and 5 eV (multipole–quadrupole); minimum quadrupole mass *m/z* 300. Data was acquired using timsControl (v6.1.5.0) and converted to .imzML format with SCiLS Lab (v2025b Pro). Slides were stored at -80°C after MALDI-MSI analysis for up to 5 days until the 10x Xenium run.

### Xenium spatial transcriptomics

Samples were analyzed based on the following demonstrated protocols, supplied by 10x Genomics. All deviations from these protocols are stated below.

1. CG000581 Rev. D: Xenium in Situ for Fresh Frozen Tissues – Fixation & Permeabilization
2. CG000582 Rev. H: Xenium in Situ Gene Expression
3. CG000584 Rev. G or K: Xenium Analyzer

#### Fixation & Permeabilization

Post-MALDI slides were retrieved from -80 °C storage and thawed on a thermocycler at 37°C for 1 min. Methanol was pre-chilled at -20°C and slides were washed through the immersion in the following solutions: twice 3 min in 100% methanol, 1 min in 70% methanol, and thrice 1 min in 1x PBS. Cells were subsequently fixed by immersion into 3.7% formaldehyde for 15 min at RT, followed by three 1 min 1x PBS washes, and finally permeabilized by immersing in 100% methanol for 1h at -20°C. After permeabilization, slides were washed twice in 1x PBS, inserted into Xenium cassettes, and PBS-T was added to the cassette.

#### Probe Hybridization, Ligation & Amplification

PBS-T was removed from the cassette before the addition of Probe hybridization mix and a subsequent incubation for 18h in a thermocyler at 50 °C. Cells were washed twice with PBS-T before the addition of post hybridization wash buffer (10x Genomics, PN-2000395) and incubation for 30 min in a thermocycler at 37°C. Slides were washed thrice with PBS-T and Ligation mix was added before incubation for 2h in a thermocycler at 37°C. Amplification master mix was added after three PBS-T washes, and the slides were incubated for another 2h in a thermocycler at 30°C.

#### Nuclei staining

Slides were retrieved from the thermocycler and washed thrice with TE buffer (Fisher Scientific, BP24731), followed by three PBS-T washes. Nuclei were stained by the addition of nuclei staining buffer (10x Genomics, PN-2000762) for 1 min. Lastly the cells were washed four times with PBS-T. Reduction and autofluorescence quenching steps were not performed.

#### Xenium Analyzer

Samples were analyzed on a Xenium analyzer instrument (10x Genomics, PN-1000481) using one of the following combinations: CG000584 Rev. G, instrument software version 3.3.0.1 and Analysis version xenium-3.3.0.1, or CG000584 Rev. K, instrument software version 4.0.1.4 and Analysis version xenium-4.0.1.0.

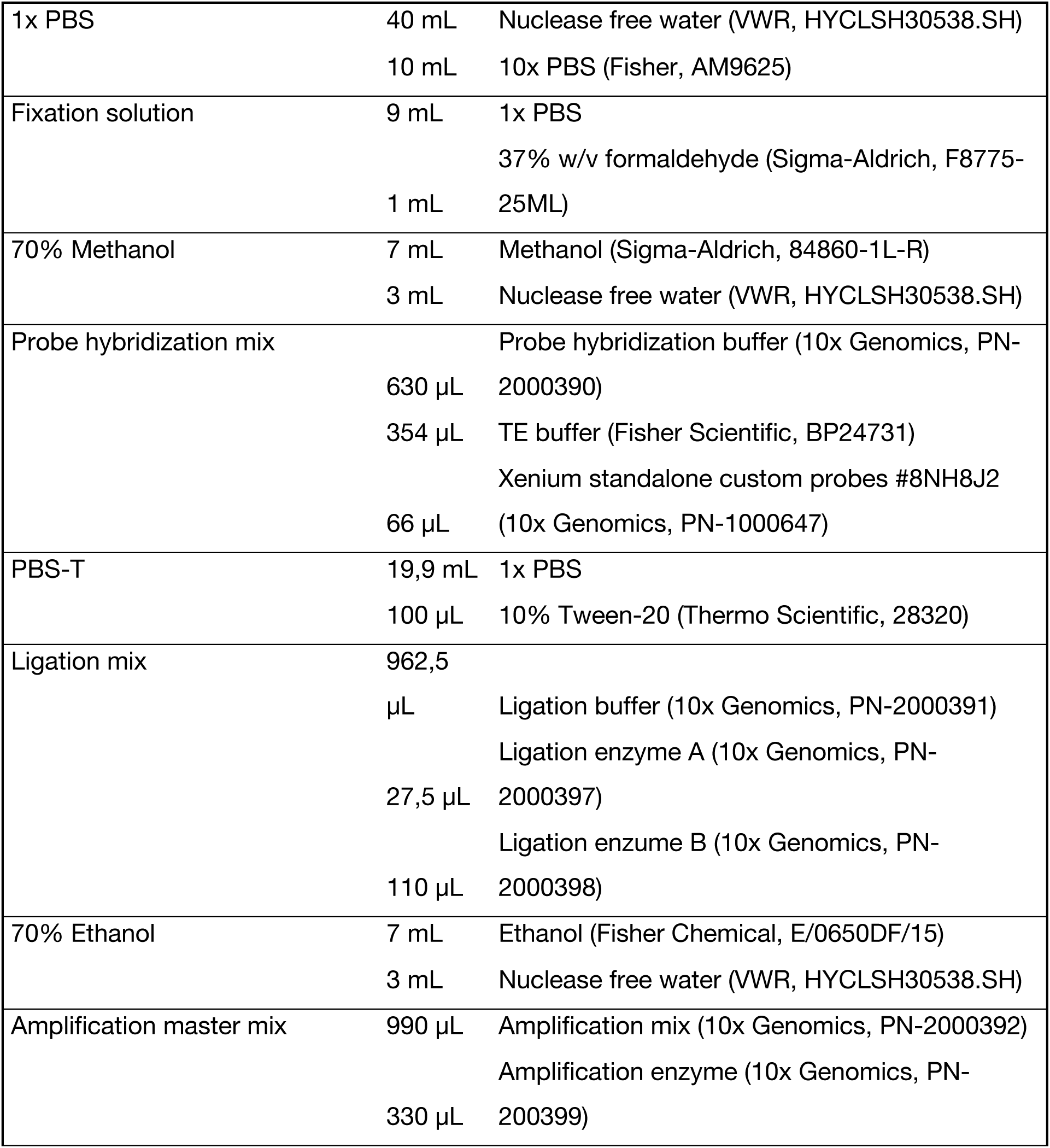

### Preparation of cells for scRNA-sequencing

In parallel with seeding cells on Xenium slides, cells were also seeded on standard glass microscope slides for generation of single-cell RNA and gRNA libraries. Glass slides were coated with fibronectin (Sigma-Aldrich, F0895) for 1 h at room temperature and allowed to dry. After three washes with DPBS, cells were seeded at the same density and at the same time as for the Xenium slides. The following day, cells were collected using 0.25% trypsin and processed according to the 10x Genomics Single Cell Protocols, Cell Preparation Guide (Rev. C). Briefly, cells were centrifuged at 300 rcf for 5 min and resuspended in 1x DPBS containing 0.04% (w/v) BSA. Suspensions were filtered through 40 µm Flowmi® cell strainers (Sigma-Aldrich, BAH136800040) and loaded onto the 10x Genomics Chromium X instrument for single-cell partitioning.

Single-cell RNA-seq libraries were prepared using the Chromium Single Cell 3′ Reagent Kits (v3.1 Chemistry, Dual Index, CG000315 Rev E) according to the manufacturer’s instructions. gRNA libraries were generated from cDNA. For sgRNA enrichment and indexing, 10% of the cDNA was used as input in a PCR reaction containing 1× KAPA HiFi HotStart ReadyMix (Roche, 7959079001), 0.5 µM sgRNA enrichment primer (5′-AATGATACGGCGACCACCGAGATCTACACXXXXXXXXACACTCTTTCCCTACACGACGCTCTTCCGATCT-3′), and 0.5 µM sgRNA primer (5′-AATGATACGGCGACCACXXXXXXXXACACTCTTTCCCTACACGACGCTCTTCCGATCT-3′). PCR products were purified using double-sided AMPure XP bead purification (Analis, BEC-A63881).

### siRNA transfection for bulk comparisons

To benchmark the LipoGrid measurements, we performed bulk lipidomics following siRNA-mediated knockdown of eight selected target genes (LPCAT3, ACBD5, BRD4, PNPLA2, TTF1, PCYT2, ELOVL5, and CHD4). RNA interference was performed by transfecting siRNA oligos using Lipofectamine RNAiMAX (Thermo Fisher Scientific, 13778) according to the manufacturer’s instructions. The following siRNA sequences were used: BRD4 (Horizon Discovery, ON-TARGETplus, J-004937-06-0002), PCYT2 (Horizon Discovery, ON-TARGETplus, J-009114-09-0002), PNPLA2 (Horizon Discovery, ON-TARGETplus, J-009003-09-0002), ELOVL5 (Horizon Discovery, ON-TARGETplus, J-009260-09-0002), LPCAT3 (Horizon Discovery, ON-TARGETplus, J-010273-09-0002), ACBD5 (Horizon Discovery, ON-TARGETplus, J-016183-18-0002), TTF1 (Horizon Discovery, ON-TARGETplus, J-012378-05-0002), and CHD4 (Thermo Fisher Scientific, Silencer Select, s2984). The ON-TARGETplus Non-targeting Pool (Horizon Discovery, D-001810-10) was used as a negative control.

Cells were transfected a second time after 4 days. Three days after the second transfection, two cell pellets were collected per condition: one for RT-qPCR analysis and one for bulk lipidomics.

#### RT qPCR for bulk comparisons

Total RNA was extracted using the Monarch RNA Extraction Kit (Bioke, T2010S). For cDNA synthesis, 100 ng total RNA was reverse transcribed using Maxima Reverse Transcriptase (Thermo Fisher Scientific, EP0752). Quantitative PCR was performed on a QuantStudio 6 Pro Real Time PCR System using 1 µL cDNA and PowerUp SYBR Green Master Mix (Thermo Fisher Scientific, A25742). Gene expression was normalized to the housekeeping genes of *ACTB* and *RPL13A*. Knockdown efficiency was calculated relative to cells transfected with the non-targeting siRNA pool.

### Bulk lipidomics

Lipid extraction, LC-MS/MS analysis, and data processing were performed as described previously^28^. Briefly, cells were homogenized using a handheld sonicator. An aliquot corresponding to 10 μg DNA was subjected to lipid extraction in the presence of HCl/methanol, chloroform, BHT, and UltimateSPLASH™ ONE and SphingoSPLASH™ I internal standard mixes. The organic phase was collected, evaporated under vacuum at room temperature, and lipid pellets were stored at -20°C under argon until analysis.

Prior to mass spectrometry, lipid pellets were reconstituted in ethanol and analyzed by LC-ESI/MS/MS using a Nexera X2 UHPLC system coupled to a 6500+ QTRAP mass spectrometer. Lipid species were separated on an XBridge amide column and quantified by scheduled multiple reaction monitoring in positive or negative ion mode. The analysis covered major glycerolipid, glycerophospholipid, sphingolipid, and cholesteryl ester species, including TAG, DAG, PC, LPC, PE, LPE, PG, PI, PS, SM, CE, CER, DCER, HCER, and LCER.

Peak integration was performed using MultiQuant™ software. Lipid signals were corrected for isotopic contributions and quantified relative to internal standards, in accordance with Lipidomics Standards Initiative guidelines for level 2 type quantification.

#### Computational Analyses

##### MALDI–MSI data pre-processing

Raw MSI data were acquired as centroided spectra at each spatial location (spot), where each spot is represented by an irregular set of detected *m/z* values with corresponding relative intensities. This variable-length representation is well suited for data storage but is not directly compatible with downstream multi-sample modeling, where each spot must be expressed on a common set of features. The overall goal is to create a dense feature matrix in which rows correspond to spots and columns correspond to a shared set of *m/z* features across all samples. We therefore applied a preprocessing workflow based on the FOCUS^22^ framework to convert the raw spectra into consistent, sample-comparable matrices.

Briefly, the workflow (i) corrects systematic mass shifts to improve cross-sample alignment; (ii) identifies tissue-containing (foreground) spots to avoid defining features from matrix-dominated background signals; and (iii) estimates a global *m/z* feature backbone using foreground spectra pooled across samples, ensuring that the final feature set is determined in a data-driven and unbiased manner rather than tuned to any single sample. Finally, all spots (foreground and background) are interpolated onto this common backbone, yielding dense intensity matrix of size nspots × nfeatures. To ensure a stable and reproducible downstream feature space, this preprocessing was applied before batch correction and statistical modelling, and all subsequent analyses were performed on either the resulting feature matrix or its batch-corrected counterpart.

The resulting matrix provides a standardized representation of MSI signal across the 4 LipoGrid runs and served as input for subsequent bioinformatics analyses. After pre-processing, the final MSI matrix comprised nspots = 1,037,286 spots and nfeatures = 7,420 features (5,472 in positive and 1,948 in negative mode).

##### MALDI-MSI alignment with Xenium DAPI signals

To align the MALDI-MSI and Xenium spatial transcriptomics to single-cell accuracy, we first need to identify spots in the MALDI-MSI datasets that likely contain a cell. For this, we applied a mask on the MALDI-MSI spots, retaining spots whose *m/z* intensities were enriched for annotated lipid species. This way we obtained a rough grid pattern of spots that likely contained a cell that was then used to align with the nuclear DAPI staining obtained from the Xenium runs. Nuclear coordinates for each cell were transferred to the MSI spot grid. This registration step was performed by interactively overlaying the MSI spot layout on the corresponding fluorescence DAPI image and manually refining the alignment. To achieve this, we relied on the alignment tool provided by the FOCUS framework^22^. The mapping from MSI coordinates to image pixel coordinates was modelled using a 2D similarity transformation (uniform scaling, rotation, and translation). After estimating this transformation, each MSI spot was projected into the Xenium image coordinate system and vice versa.

The projected Xenium cell coordinates seldom coincide exactly with discrete MSI pixels and usually intersects multiple neighboring spots. Hence, we employed a Gaussian-weighted interpolation filter on the closest 4 MALDI-MSI spots for each cell. This filter distributes each MSI vector proportionally across the overlapping Xenium spot area, thereby preserving spatial fidelity while accommodating sub-pixel alignment. Across the 4 slides, a total of 79,521 cells were aligned, and an intensity profile for the synchronized 7,420 *m/z* values was calculated for each cell.

#### DAPI segmentation on Xenium output and gRNA count-matrix creation

Owing to the simplicity of the staining due to the grid-based experimental design, we noticed that our own optimized segmentation pipeline recovered more cells and transcripts compared to the standard 10x Xenium pipeline. Nuclei were segmented using traditional thresholding followed by watershed refinement. More specifically, the staining was first blurred by a gaussian blur of *sigma*=1 (Implementation of *skimage.filters.gaussian*, version==0.24.0). After this, foreground was split from background by calculating an otsu’s threshold (*skimage.filters.threshold_otsu*) based on a subsampled version of the staining to avoid perturbance by technical artefacts. A distance transform (*scipy.ndi.distance_transform_edt*, version==1.13.1) was applied to the resulting mask, which was then used to calculate local maxima in the mask (*skimage.feature.peak_local_max*, min_distance=5). Final nuclei segmentation is obtained by applying a watershed algorithm on the distance image using the local max peaks as seeds (*skimage.segmentation.watershed*). The segmentation boundaries were then expanded by 30 pixels. While doing so, segmented cells that were touching after expansion were logged. A graph of touching cells was constructed using *networkx* (version==3.2.1). Detected spots of the Xenium experiment were then assigned to cells based on which segmentation mask they were included in. The resulting data object was saved to an *AnnData* (version == 0.10.8) array, compatible with standard Spatial Transcriptomics workflows.

##### gRNA assignment

A cell was assigned a single gRNA (and corresponding target gene) if that gRNA was detected ≥8 times, was ≥1.5-fold more abundant than the next most frequent gRNA, and accounted for >25% of total gRNA counts in that cell. This resulted for the 79,521 aligned cells in 57,743 cells with a dominant single gRNA, 15,045 cells expressing multiple gRNAs, and 6,733 cells with low or no detectable gRNA expression (Figure 1C).

##### Enriching for biological signal in *m/z* spectra

A substantial amount of the 7,420 detected *m/z* values does not come from actual biomolecules but rather originates from the matrix used in MALDI-MSI or other non-biological sources. An advantage of having the cells spaced in a micro-grid is that we also know which locations are devoid of cells. We used this knowledge to calculate an average background signal for each *m/z* value. Only *m/z* values whose average intensity is at least 1.8 times higher in the cells compared to the background are retained, reducing to number of *m/z* values to 2,580 (2,253 in positive and 327 in negative mode) (Figure 1E).

Using this set of biologically enriched *m/z* values, we calculated for each cell its total intensity profile. Similar to standard quality-control procedures in single-cell RNA-seq, we excluded cells with abnormally low (mean - 1.25 × SD) or high (mean + 2.5 × SD) total signal intensity in either positive or negative ion mode, as these likely represented low-quality measurements. This way we filtered out 8,877 low quality cells, ending up with 70,644 cells with an informative MSI profile in positive ion mode and 47,835 in negative mode.

Positive- and negative-ion mode data were analyzed separately and on these different cell sets throughout the paper. Positive mode was acquired in all four runs while negative mode was acquired only in runs 3 and 4; all negative-mode steps, TIC normalisation, control-anchored run correction, feature filtering and differential testing, were therefore restricted to the n = 47,835 cells from those two runs, using the intergenic-control cells of the same two runs as reference. No negative-mode values were imputed for runs 1 and 2.

##### Internal normalization of *m/z* signal

We normalized the intensity profile of each cell making use of the internal control cells (gRNAs targeting intergenic regions) in each of the 4 MALDI-MSI runs and per ion mode. First, to correct for cell-to-cell variation in total lipid signal, analogous to read-depth normalization in single-cell RNA-seq, we applied total ion current (TIC) normalization. For each cell, the total intensity was computed as the sum of all intensity values across the 2,253 *m/z* features in positive mode or the 327 *m/z* features in negative mode. The mean total intensity across cells was then calculated for each mode, and a per-cell scaling factor was defined as the ratio of that cell’s total intensity to the mean total intensity. Each cell’s intensity values were divided by its scaling factor to yield the normalized profile. Secondly, to remove technical variation between the four MALDI-MSI runs, we used the internal control cells (gRNAs targeting intergenic regions), present in every run and expected to share a common average lipid composition, as an anchor. This step followed per-cell TIC normalization and was performed separately for positive and negative mode. For each run we computed a reference control profile by averaging the normalized intensities of its control cells (one value per *m/z* feature; 2,253 in positive, 327 in negative mode), and averaged these across runs to obtain a grand-mean control profile. Per-run, per-feature correction factors were defined as the ratio of the grand-mean to that run’s control intensity (with a pseudocount of 1 added to numerator and denominator to avoid division by zero and damp low-intensity features), clipped to [0.4, 2.5] to prevent extreme scaling. Each feature’s intensities in all cells of a run, control and perturbed alike, were then multiplied by the corresponding correction factor, aligning the four runs to a shared, control-defined intensity scale (See UMAPs before and after Figure S3).

##### Generation of final cells over *m/z* intensity matrix

Finally, we further removed low-intensity signals while preserving biologically relevant signals, including species potentially induced by specific knockouts. For this, we grouped the cell based on their gRNA target gene and calculated an average intensity profile for each of them. An *m/z* value was retained if for at least one target gene, its average intensity passed the cutoff. This resulted in a final feature set of 1,160 *m/z* values in positive and 217 in negative mode for 51,507 cells with a single dominant gRNA expressed.

##### Evaluating the impact of each gene knockout on the *m/z* intensity profile

We continued the analysis using the 51,507 high-quality cells that expressed a single dominant gRNA and grouped them according to their target. In total, 153 target genes and 20 gRNAs targeting intergenic control regions were initially selected (Table S1). For downstream analyses, we retained only those target genes whose corresponding gRNAs were detected in at least 120 cells, resulting in 143 target genes included for further analysis (Figure. 1D).

The impact of each gene knockout on the *m/z* intensity profile was calculated by grouping the cells by target gene (or intergenic control) and averaging their intensity profile for each of the 1,377 *m/z* features. The log_2_ fold change for each *m/z* feature was calculated over the internal controls as follows: log_2_((avg intensity target gene + 1)/(avg intensity intergenic controls +1)).

The impact of each gene knockout was visualized using the log_2_FC values per gene over all *m/z* features in a heatmap (Figure 2A). Log_2_FC outliers were clipped to [-1.5,1.5] for visualization. Target genes and *m/z* features were clustered (using AgglomerativeClustering, linkage=Ward) and the *m/z* features were further ordered using hierarchical clustering. Additionally, the log_2_ total average intensity for each *m/z* feature over all cells is plotted underneath the heatmap. Followed by the log_2_FC of this average total intensity in the cells over the average total intensity in the surrounding matrix. The final bar represents whether this *m/z* feature originated from the positive or negative ion mode.

The (dis)similarity in *m/z* intensity profile for each gene knockout is visualized using Uniform Manifold Approximation and Projection (UMAP) on the 1,377 *m/z* features and 10 neighbors setting, the TF mentioned in the main text are colored in red and the MGATs in green.

##### Gene Ontology enrichment analysis

Comparative Gene Ontology enrichment analysis on the 4 target gene clusters (Fig 2A,) was performed using g:Profiler^24^ using multi query with standard settings. The highest scoring (p-value based) ontologies that displayed specific enrichments were selected for display. GeneRatio displays the ratio of genes in the cluster that overlap with the ontology term.

##### Linking *m/z* values to annotated lipids

We used a by lipid experts curated stringent *m/z* to ionized lipid species list for both positive and negative ion mode MALDI-MSI. An *m/z* feature was linked to an ionized lipid specie if its value was within the 10-ppm machine mass tolerance according to the formula;

| mz_feature – mz_lipid | / mz_feature =< 10 x 10^-6^

We matched 182 of the 1,377 filtered *m/z* features to an ionized lipid specie. After collapsing multiple ionization states of the same molecule, this yielded 158 unique lipid species (Table S4).

##### Statistical analysis of gene knockout impact on measured lipidome

We quantified for each of the 158 lipid species detected in our screen their abundance changes due to each gene knockout. For this we calculated log_2_ fold change for each lipid as follows: log_2_((avg lipid intensity target gene + 1)/(avg lipid intensity intergenic controls +1)). To estimate the significance, a p-value was calculated using the non-parametric statistical Mann-Whitney U test using all cell with a specific gene knockout versus the cells with gRNAs targeting intergenic regions as controls.

##### Comparison to bulk lipidomics

From the bulk lipidomics dataset, we used the quantile-normalized abundance of each detected lipid species, expressed as nmol per mg DNA. Because bulk lipidomics resolves individual fatty acyl chain compositions, whereas MALDI-MSI reports lipids at the sum-composition level, lipid species differing only in fatty acyl chain composition were aggregated into their corresponding sum notation to enable direct comparison between the two platforms. Using six control replicates and three siRNA knockdown replicates for each target gene, we calculated log_2_fold changes and assessed statistical significance using a two-sample independent *t*-test.

LipoGrid results were compared directly with bulk lipidomics by considering lipid species detected by both platforms. Of the 158 lipid species identified by LipoGrid, 97 had a one-to-one match in the bulk lipidomics dataset. For each of the eight perturbations (LPCAT3, ACBD5, BRD4, PNPLA2, TTF1, PCYT2, ELOVL5, and CHD4), we selected lipid species that changed significantly (P < 0.05) in both the bulk lipidomics and MALDI-MSI datasets. This yielded 57 lipid species–gene perturbation pairs for comparison of the log_2_ fold changes between the two platforms (Figure S5). Concordance between the datasets was assessed using the Pearson correlation coefficient.

##### Data visualization using boxplots and volcano plots

###### Boxplot generation

Boxplots summarizing grouped datapoints were generated in Python using matplotlib. For each group the mean log₂ fold change of the lipids or *m/z*s values for the given gene knockout over controls was plotted. Boxes span the interquartile range with the median indicated by a horizontal line, and whiskers extend to the most extreme non-outlier values. Individual data points were overlaid as jittered scatter points, colored by cluster, and a dashed horizontal reference line was drawn at zero.

###### Volcano plot generation

Volcano plots for individual gene knockouts or lipid species were generated in Python using matplotlib. For a given gene or lipid, the log₂ fold change (gRNA relative to intergenic control gRNA) and associated p-value were plotted for all lipid species or gene knockouts respectively. P-values were capped at a lower bound of 1 × 10⁻^10^ to limit the vertical extent of highly significant points and plotted as -log₁₀(p-value) against log₂ fold change. Significant (P < 0.01) genes/lipids were named; colored blue when depleted (log₂FC < 0) and red when enriched (log₂FC > 0).

##### CROP-seq pre-processing

To map the single-cell RNA-seq read we ran CellRanger from 10x genomics. To map the CROP-seq reads (single cell RNA and gRNAs) we first made a custom reference that included all possible gRNAs together with the human CRCh38 transcriptome. For this we generated a fasta file and gtf file of the gRNAs transcripts and concatenated these files to the Homo_sapiens.GRCh38 assembly fasta and gtf files. We then made a new reference using:

cellranger mkref with the following parameters:

- -genome=AddedOligoPool_GRCh38
- -fasta=Homo_sapiens.GRCh38_gRNAoligo_assembly.fa
- -genes=Homo_sapiens.GRCh38.93_gRNAoligo.gtf
- -ref-version=3.0.0

cellranger mkgtf Homo_sapiens.GRCh38.93.oligo.gtf Homo_sapiens.GRCh38.93.oligos.filtered.gtf \

- -attribute=gene_biotype:protein_coding \
- -attribute=gene_biotype:lincRNA \
- -attribute=gene_biotype:antisense \
- -attribute=gene_biotype:IG_LV_gene \
- -attribute=gene_biotype:IG_V_gene \
- -attribute=gene_biotype:IG_V_pseudogene \
- -attribute=gene_biotype:IG_D_gene \
- -attribute=gene_biotype:IG_J_gene \
- -attribute=gene_biotype:IG_J_pseudogene \
- -attribute=gene_biotype:IG_C_gene \
- -attribute=gene_biotype:IG_C_pseudogene \
- -attribute=gene_biotype:TR_V_gene \
- -attribute=gene_biotype:TR_V_pseudogene \
- -attribute=gene_biotype:TR_D_gene \
- -attribute=gene_biotype:TR_J_gene \
- -attribute=gene_biotype:TR_J_pseudogene \
- -attribute=gene_biotype:TR_C_gene

Mapping of the CROP-seq runs was performed using CellRanger/8.0.1 with the following parameters for both the sc-RNA-seq library and the gRNA enriched library:

cellranger count --id=Chemv3_lipogrid_CROP_extraseq transcriptome=AddedOligoPool_GRCh38/

- -chemistry SC3Pv3
- -create-bam=true

##### CROP-seq analysis

###### Single-cell RNA-seq

Reads were mapped and barcodes deconvoluted using CellRanger, yielding RNA-seq profiles for 33,617 cells. Standard quality control was performed, removing cell with low read counts and potential doublet outliers by filtering cells on total counts (8.4 < log1p total counts < 10.7), which retained 31,210 cells. These were library-size normalized and to preserve genes that might be induced by individual knockouts, a lenient expression filter was applied, retaining genes for which (gene total counts + 1)/(number of cells in which the gene was detected + 1) > 1.1. This yielded 11,506 genes for downstream analysis. *gRNAs assignment.* To quantify the gRNAs expressed in each cell, we mapped the second, gRNA-enriched library (see Preparation of cells for scRNA-sequencing), obtaining 25,086 cells from CellRanger. These were overlapped by barcode similarity with the 31,210 cells detected in the single-cell RNA-seq experiment, yielding 24,697 cells with both gRNA information and high-quality scRNA-seq data. As in the 10x Xenium analysis, we then examined the distribution of detected gRNAs per cell: 15,346 cells expressed a single dominant gRNA, 9,250 expressed multiple gRNAs, and 101 showed low or ambiguous gRNA expression. Downstream analysis was continued with the 15,346 cells carrying a dominant assigned gRNA.

###### Differential expression analysis per gene knockout

Similar to the lipidome analysis, we quantified transcriptional changes for each perturbation by comparing gene expression in cells carrying a given target gRNA with cells harboring intergenic control gRNAs. For each gene, the log₂ fold change was calculated from the mean expression in target and control cells as log₂((mean target + 1)/(mean intergenic + 1)), and statistical significance was assessed using a two-sided Mann–Whitney U test. This yielded perturbation-specific log₂ fold changes and p-values for all measured genes.

##### Gene Set Enrichment Analysis

###### Compilation of lipid-related gene sets

To enable lipid-focused enrichment analyses, we assembled a custom gene set (GMT) collection restricted to lipid-related terms drawn from multiple pathway databases: MSigDB Hallmark, Reactome, KEGG, Gene Ontology Biological Process, the broader MSigDB canonical pathway collections (C2CP, including BioCarta, PID, and WikiPathways) and GO (C5), as well as PathBank. Terms were classified as lipid-related by matching of their names against a curated list of keywords spanning lipid classes (e.g. cholesterol, sterols, phospho-, sphingo- and glycerolipids, fatty acids), lipid metabolism (e.g. β-oxidation, lipogenesis, lipolysis, the mevalonate and eicosanoid pathways), lipid transport and lipoproteins (e.g. LDL, HDL, apolipoproteins), and key lipid-associated regulators (e.g. PPAR, SREBP). To avoid redundancy across databases, term names were canonicalized by stripping database-specific prefixes and trailing identifiers and collapsing to a normalized key; entries sharing a canonical key were merged into a single gene set by taking the union of their gene members, while recording the contributing source databases. The resulting lipid-focused gene set collection was written to a GMT file for use in subsequent enrichment analyses.

###### Batch GSEA

Gene set enrichment analysis across knockouts. For each gene knockout, genes were ranked by a signed significance score, computed as sign(log₂ fold change) × - log₁₀(p-value). Pre-ranked GSEA was then run on each knockout’s ranking against the custom lipid-related gene set collection using GSEApy (gp.prerank; minimum gene set size 5, 1,000 permutations). Normalized enrichment scores (NES), nominal p-values, and FDR q-values were collected across all knockouts and assembled into term × ranking matrices. Nominal p-values were additionally corrected within each ranking using the Benjamini-Hochberg procedure. For visualization, terms were retained if they reached an adjusted p-value below 0.1 in at least one ranking (top two per ranking), and their NES values were displayed as a hierarchically clustered heatmap, with adjusted-p-value significance indicated by asterisks (* < 0.25, ** < 0.05, *** < 0.01).

##### Arrow plot generation

Arrow plots integrate pathway enrichment with lipidomic and transcriptional data across the gene knockout panel. For a given pathway term and lipid species, each knockout was plotted by its GSEA normalized enrichment score (NES, y-axis) against the lipid log_2_FC over intergenic controls (x-axis). Knockouts were colored by significance on each axis: GSEA FDR ≤ 0.1 in blue, lipid P ≤ 0.01 in orange, and additional genes involved in the pathway in grey. Directed arrows between labeled genes were overlaid from the CROP-seq RNA log_2_FC matrix, drawn from knockout gene to target whenever knockout significantly changed the target’s expression (P ≤ 0.01, |log2FC| ≥ 0.2), with color denoting up-versus downregulation and reciprocal pairs curved apart.

##### Lipid-import genes

We investigated the expression changes of all candidate lipid import genes. The original lists were; cholesterol = [LDLR, VLDLR, LRP1, LRP2, LRP4, LRP5, LRP6, LRP8, SCARB1, SCARB2, CD36, CXCL16, OLR1, MSR1, NPC1L1, SORT1, SORL1, LRPAP1, APOE, APOB, APOA1, APOA4, LPL, LIPC, LIPG and LIPA], Fatty Acids = [CD36, SLC27A1, SLC27A2, SLC27A3, SLC27A4, SLC27A5, SLC27A6, FABP1, FABP2, FABP3, FABP4, FABP5, FABP6, FABP7, CAV1 and CAV2]. We filtered out genes that were not/lowly expressed in our cell line (mean counts >= 1). This retained LDLR and LRPAP1 for cholesterol and FABP5, CAV1 and CAV2 for Fatty Acid import. 4 core enzymes were selected for each pathway and log2FC, p-value were retrieved from the general CROP-seq analysis.

##### How to use the GUI to inspect lipids/genes of interest

To enable real-time exploration of the impact of individual gene knockouts on the measured lipidome and transcriptome, we developed a user-friendly web interface, available at https://sifrimlab.org/LipoGrid/. The data can be browsed interactively: selecting a knockout gene displays its effect on both the lipidome and the transcriptome, selecting a lipid species shows all gene knockouts that affect it, and entering a gene shows how its expression changed across the knockouts.

