## Supplementary figures for "LipoGrid: A High-Throughput Multi-omics Perturbation Screen Dissects the Genetic Architecture of Lipid Metabolism"

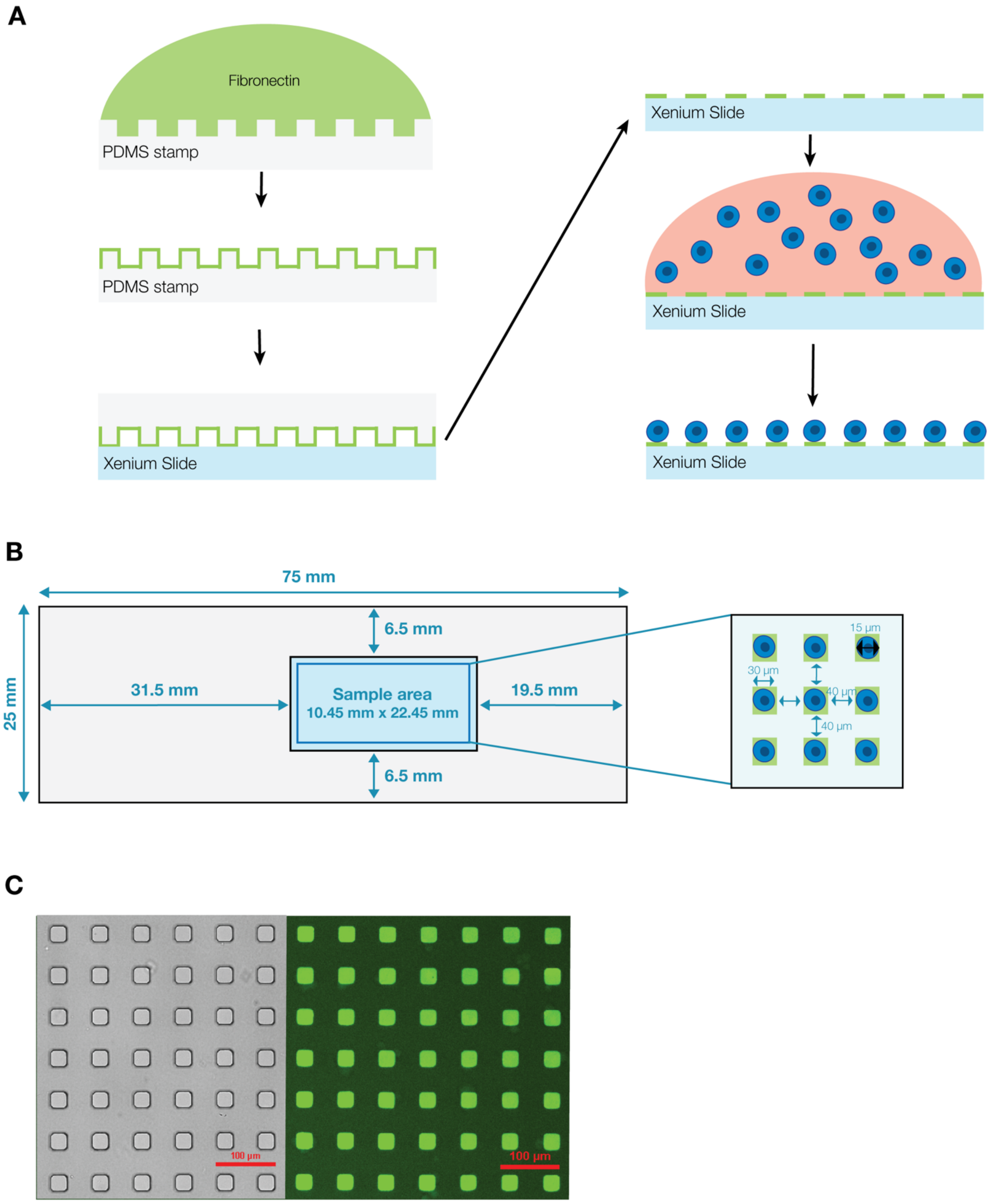


**Figure S1. Single-cell micropatterning workflow and representative results.** (A) Fibronectin is transferred from a PDMS micropillar-array stamp to the Xenium working area, after which A549 cells are seeded and selectively adhere to the fibronectin microgrid. (B) Schematic of the Xenium slide and working area, showing the dimensions of the fibronectin microgrid and A549 cells. (C) PDMS micropillar structures (30 $\mu$m) and their fibronectin-FITC micropattern grid imprints on a Xenium slide.

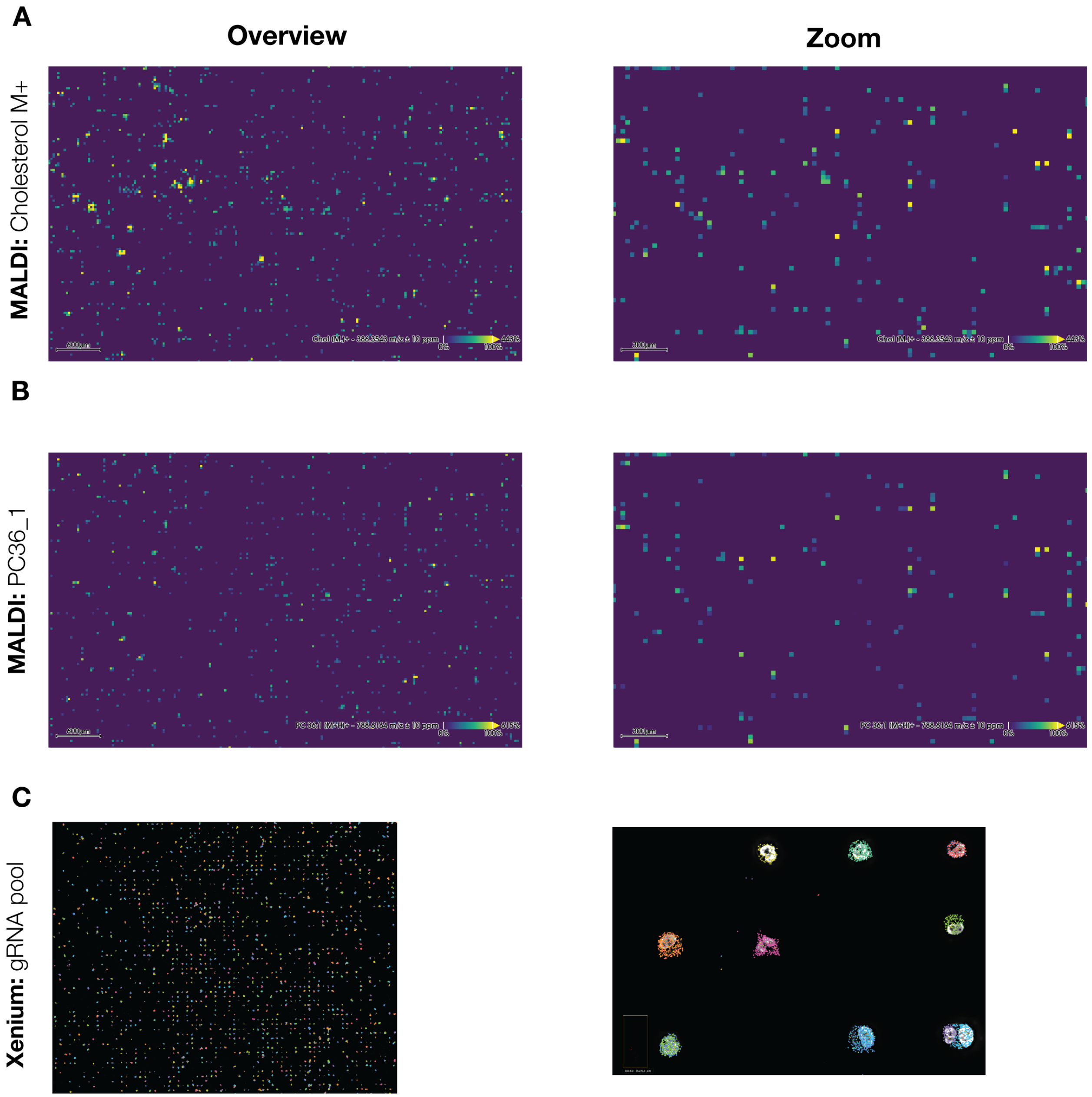


**Figure S2. MALDI-MSI screenshots and 10x Xenium gRNA visualization of LipoGrid run** (A-B) Raw ion images from MALDI-MSI, depicting the *m/z* of (A) cholesterol (M.)+, 386.3543 *m/z* and (B) PC 36:1 (M+H)+, 788.6164 *m/z*, two prominent lipids in A549 cells. (C) 10x Xenium output images, wide selection of slide and zoomed-in region, with each color representing a distinct gRNA.

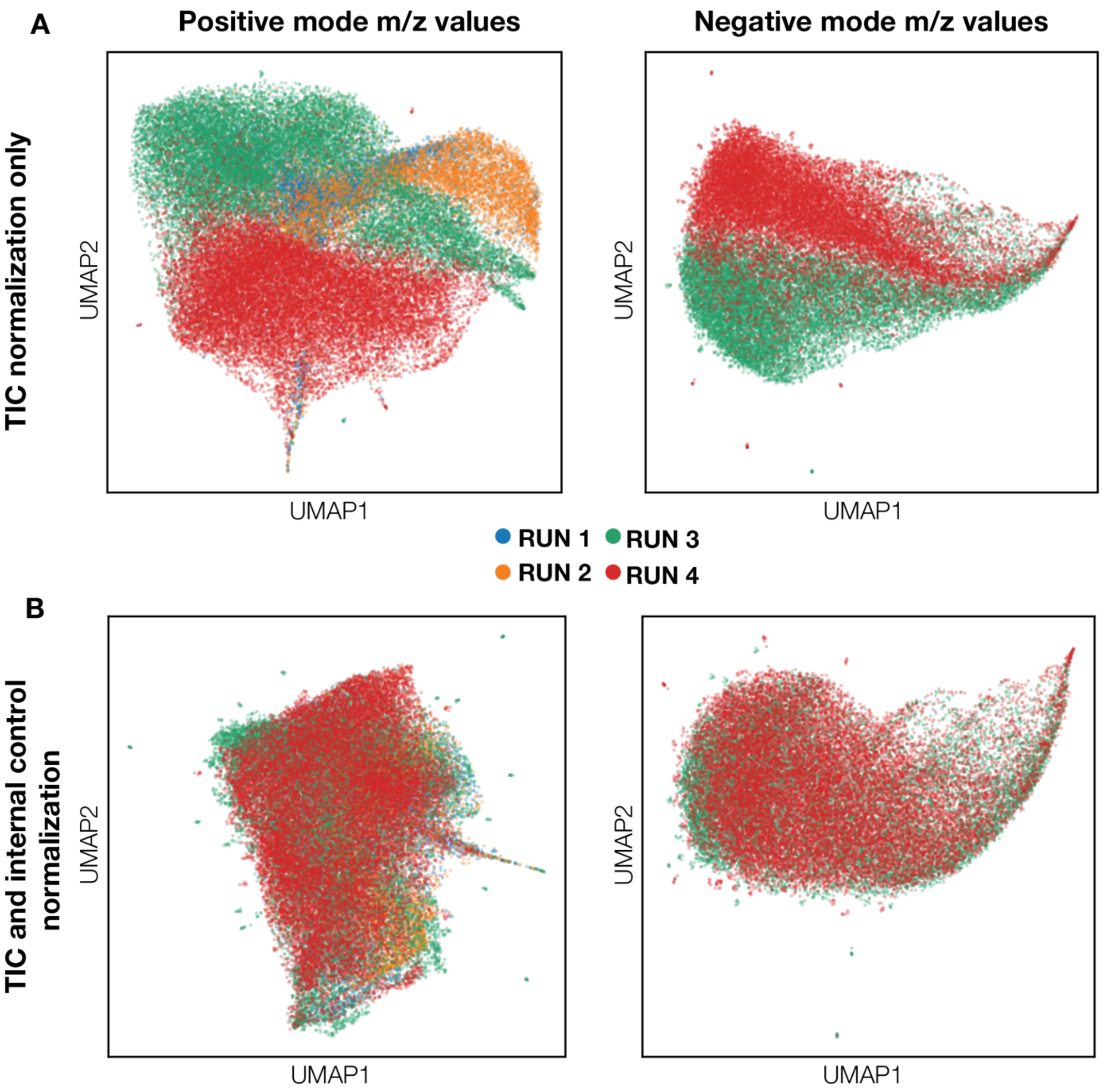


**Figure S3. Internal batch correction using intergenic targeting control cells.** (A) UMAP projection of the TIC normalized m/z intensity spectra in positive ion mode (2,253 features) and negative ion mode (327 features) for each cell, colored by run. (B) UMAP projection of the TIC and internal control normalized (see Methods) m/z intensity spectra in positive ion mode (2,253 features) and negative ion mode (327 features) for each cell, colored by run.

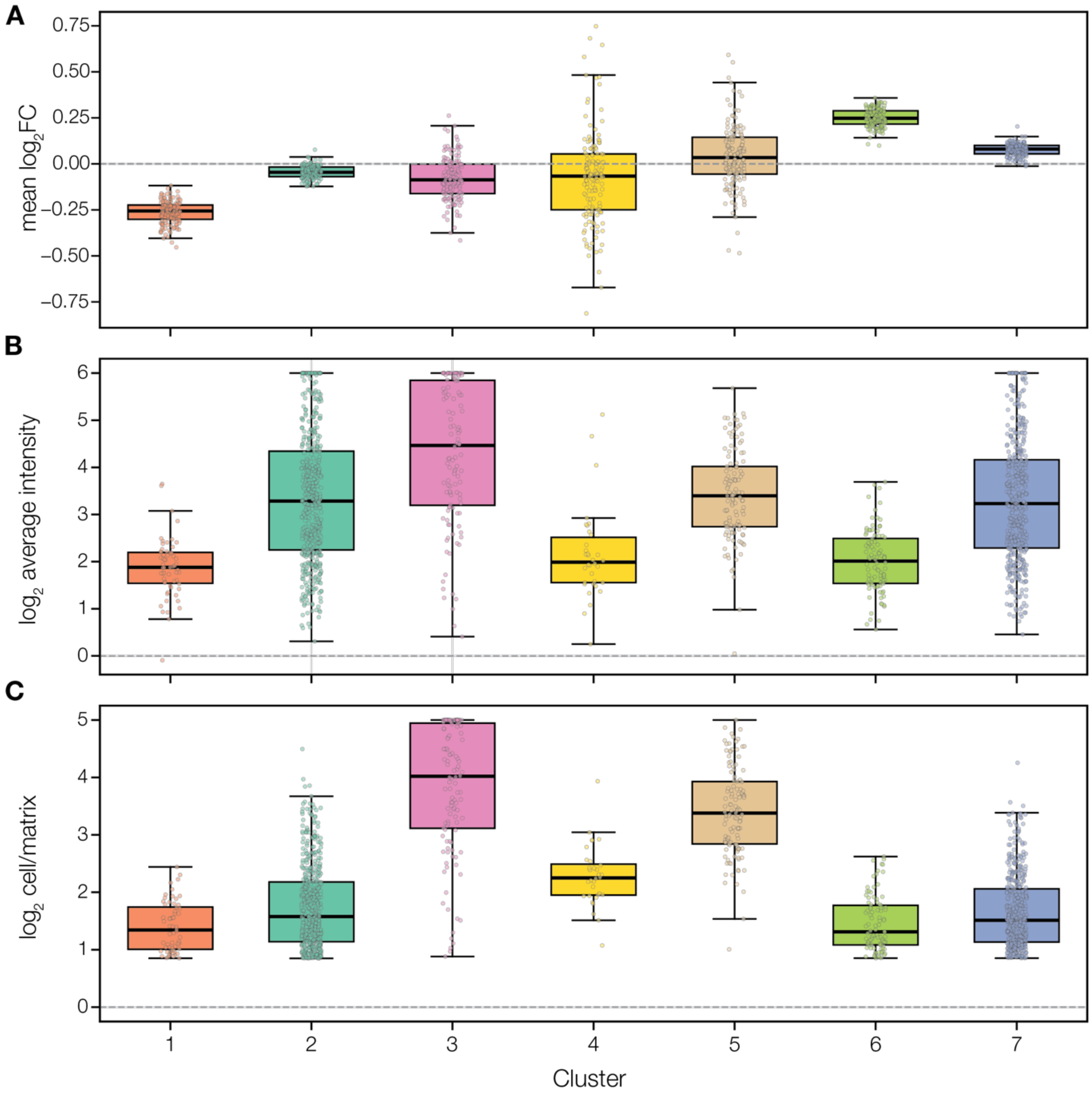


**Figure S4. Per-cluster summary of m/z feature behavior across knockouts.** (A-C) m/z features were grouped into column clusters (x-axis; numbered 1–7) by hierarchical clustering of their knockout $log_2$ fold-change profiles (heatmap 2A). Cluster sizes were n = 60, 505, 116, 34, 121, 96 and 445 m/z features for clusters 1–7, respectively (1,377 features total). For each cluster, three metrics are shown as boxplots with individual features overlaid as jittered points: (A) the mean $log_2$ fold change across all knockouts, indicating each cluster's overall perturbation-response magnitude and direction; (B) the $log_2$ average intensity, reflecting feature abundance; and (C) the $log_2$ ratio of signal in cells versus matrix background, indicating cell-specific enrichment. Boxes show the median and interquartile range (IQR), whiskers extend to 1.5× IQR, and the dashed line marks zero. Extreme outliers (beyond 3× IQR from the quartiles, Tukey "far out" rule) were removed for visualization.

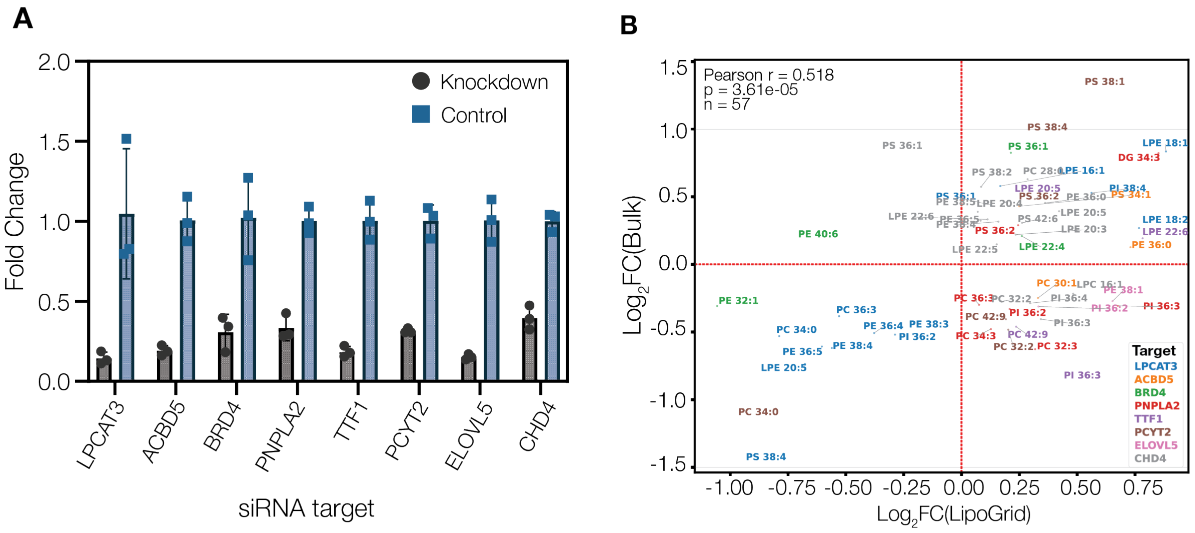


**Figure S5. Orthogonal validation of LipoGrid lipidomic effects.** (A) qPCR quantification in triplicate of siRNA-mediated knockdown for the eight target genes (LPCAT3, ACBD5, BRD4, PNPLA2, TTF1, PCYT2, ELOVL5 and CHD4), shown as paired bars (knockdown versus control) confirming reduced target expression. (B) Correlation of lipid fold changes between LipoGrid (single-cell MALDI) and bulk lipidomics. Each point is a lipid species scoring significant ($p < 0.05$) in both platforms for a given knockout, colored by target gene (legend). Axes show $\log_2$ fold change versus control in LipoGrid (x) and bulk lipidomics (y); dashed red lines mark zero on each axis. The two measurements are positively correlated (Pearson r = 0.52, p = 3.61 x $10^{-5}$, n = 57 significant lipid–knockout pairs).

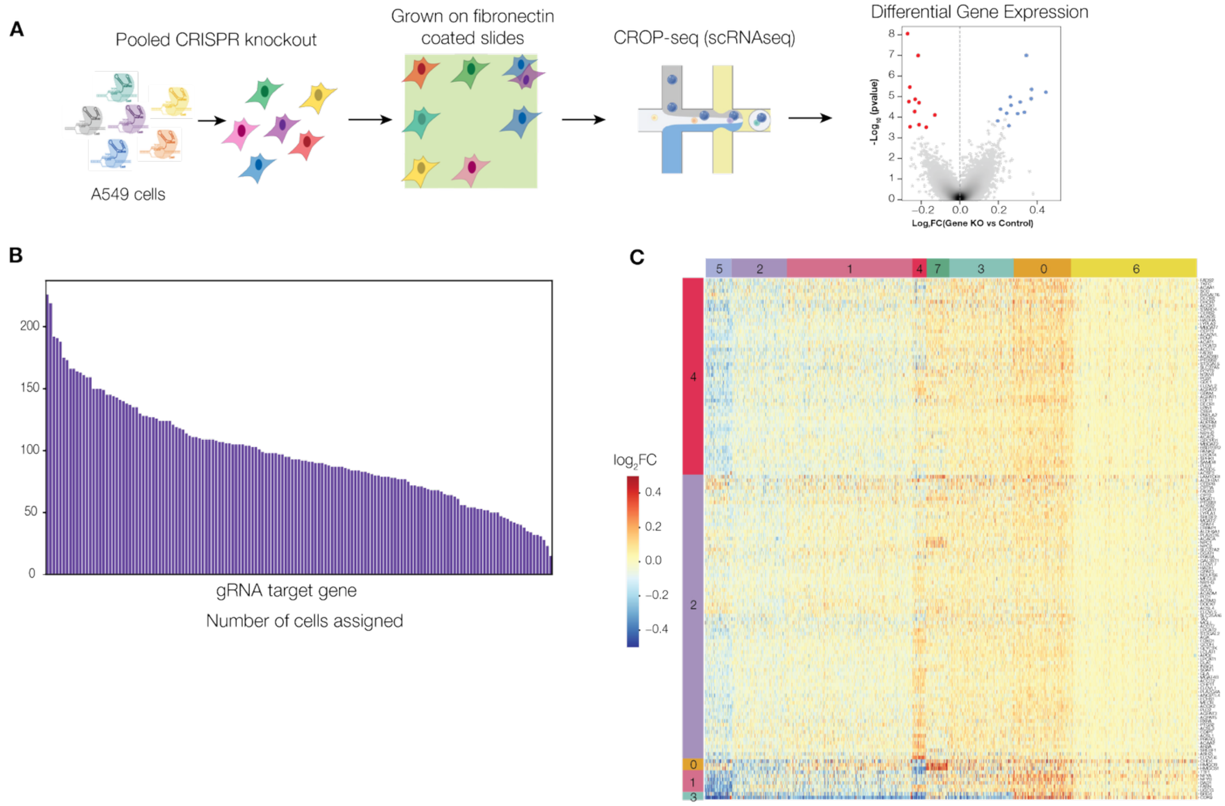


**Figure S6. Single-cell RNA-seq (CROP-seq).** (A) Schematic of parallel CROP-seq procedure to capture the changes in gene expression due to the gene knockouts. (B) Bar plot visualizing the distribution of the different genes targeted by gRNAs. (C) Heatmap showing the $log_2$ fold change in mean expression for each variable gene (FDR < 0.05 for at least one knockout, 1,674 genes) following gene knockout relative to intergenic controls across the 143 target genes (rows). Rows and columns are ordered using agglomerative hierarchical clustering.

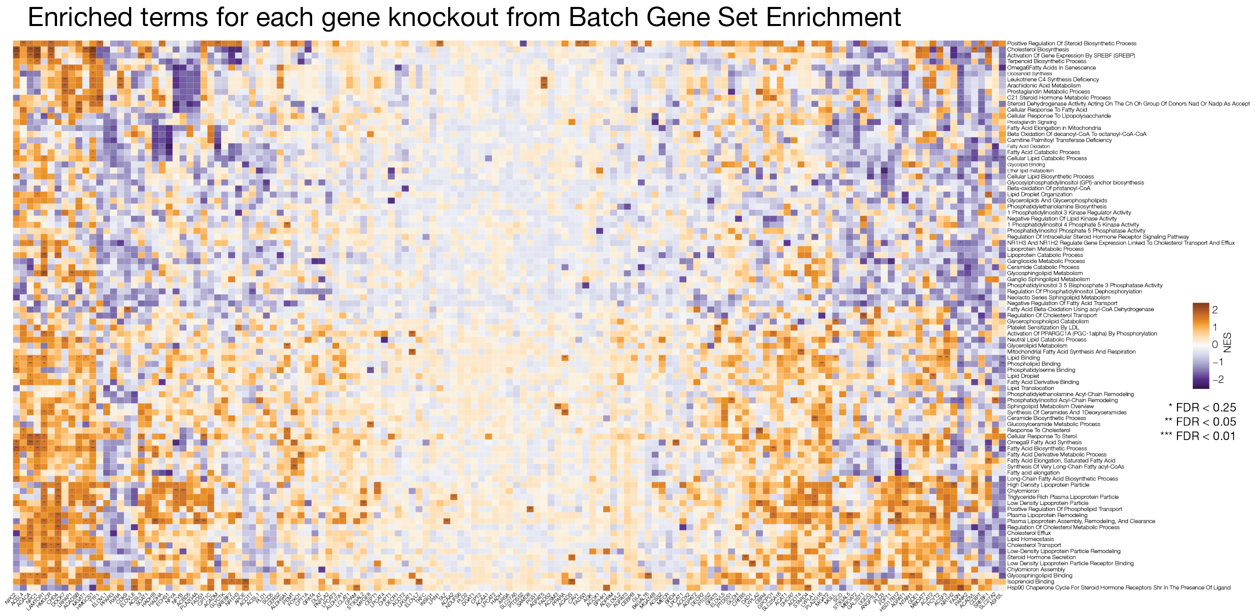


**Figure S7. Gene-set enrichment across all knockouts.** Clustered heatmap of gene-set enrichment analysis (GSEA) normalized enrichment scores (NES) for each knockout (columns) against the tested gene sets (rows). Color encodes NES (purple, positive/enriched; orange, negative/depleted). Rows and columns were hierarchically clustered. Asterisks denote statistical significance (*** < 0.01, ** < 0.05, * < 0.25) based on the GSEA FDR q-value. Only gene sets passing the significance/selection filter in at least one knockout are shown.
